# *LabGym*: quantification of user-defined animal behaviors using learning-based holistic assessment

**DOI:** 10.1101/2022.02.17.480911

**Authors:** Yujia Hu, Carrie R. Ferrario, Alexander D. Maitland, Rita B. Ionides, Anjesh Ghimire, Brendon Watson, Kenichi Iwasaki, Hope White, Yitao Xi, Jie Zhou, Bing Ye

## Abstract

Quantifying animal behavior is important for many branches of biological research. Current computational tools for behavioral quantification typically rely on a few pre-defined, simplified features to identify a behavior. However, such an approach restricts the information used and the tool’s applicability to a limited range of behavior types or species. Here we report a new tool, *LabGym*, for quantifying animal behaviors without such limitations. Combining a novel approach for effective evaluation of animal motion with customizable convolutional recurrent networks for capturing spatiotemporal details, *LabGym* provides holistic behavioral assessment and accurately identify user-defined animal behaviors without restrictions on behavior types or animal species. It then provides quantitative measurements of each behavior, which quantify the behavior intensity and the body kinematics during the behavior. *LabGym* requires neither any intermediate step for processing features that causes information loss nor programming knowledge from users for post-hoc analysis. It tracks multiple animals simultaneously in various experimental settings for high-throughput and versatile analysis. It also provides users a way to generate visualizable behavioral datasets that are valuable resources for the research community. We demonstrate its efficacy in capturing subtle behavioral changes in animals ranging from soft-bodied invertebrates to mammals.

## INTRODUCTION

Quantitative measurements of animal behavior are important for many branches of biology and biomedical research (Altimus et al., 2020; Datta et al., 2019; Egnor and Branson, 2016; Pereira et al., 2020). Identifying a behavior is a prerequisite to quantifying it. Current computational tools for behavioral quantification largely rely on tracking a set of pre-defined, simplified features to identify a behavior (Berman et al., 2014; Denisov et al., 2013; Hsu and Yttri, 2021; Kabra et al., 2013; Klibaite et al., 2017; Klibaite and Shaevitz, 2020; Nilsson et al., 2020; Pereira et al., 2019; Segalin et al., 2021; Wiltschko et al., 2015). For example, JAABA uses pre-defined features (e.g., instantaneous speed, positions, and areas) that are derived from the outlines of the animals’ bodies for identifying a set of behaviors in mice and flies (Kabra *et al.*, 2013). Similarly, B-SOiD uses body part positions computed by DeepLabCut (Mathis et al., 2018) to generate speed, angles, and body lengths for identifying animal behaviors (Hsu and Yttri, 2021). MARS uses the postural features of two mice to classify social behaviors in mice (Segalin *et al.*, 2021). MotionMapper also uses postural features of adult *Drosophila* to identify behaviors in flies (Berman *et al.*, 2014). In *Drosophila* larvae, SALAM/LARA evaluates several specific contour features for behavior detection (Denisov *et al.*, 2013).

While tools that use pre-defined, simplified features have greatly advanced behavioral analyses, they have limitations. Firstly, the features need to be defined *a priori* for identifying certain behavior types, which may fail to generalize to other behavior types. For example, motion speed can be used to identify locomotion but is useless for distinguishing facial grooming from body grooming during which the animal is immobile. Animals can also display new behaviors in response to experimental manipulations, such as drug exposure (Ellinwood and Balster, 1974; Ferrario et al., 2005). Such qualitatively different behaviors might not be easily captured by pre-defined features. Moreover, the same behavior may be recorded from different camera angles. A tool with features that are pre-defined for side views might not work for videos captured in top-view. Secondly, feature-based approaches assume that only the features selected by the designer are key to identifying a given behavior. However, the limited number of pre-defined features typically underrepresent the complete information needed to capture the complexity of animal behavior; and defining a large set of features that encompass every aspect of a behavior *a priori* is not only computationally inefficient but also might be impossible. Finally, a feature is typically a simplified summary of one aspect of the behavior. Information loss might occur in processing the feature, which makes the behavior identification less accurate. For example, speed derived from the distance of the animals’ centers of mass is a common feature that is used in feature-based tools, but the information of animal body shape is lost in such simplification.

Recent advancements in deep learning offer new approaches for behavior analysis. For example, DeepLabCut and LEAP use deep convolutional neural networks (Fukushima, 1980; LeCun and Bengio, 1995) to enable accurate tracking of animal body parts for pose estimation (Mathis *et al.*, 2018; Pereira *et al.*, 2019). These tools track body positional in space across time, but on their own do not identify or quantify any behavior. As a result, additional post-hoc tools, such as B-SOiD (Hsu and Yttri, 2021), are required to identify and quantify a behavior based on the outputs of the tracking tools. Other deep-learning-based tools such as DeepEthogram (Bohnslav et al., 2021) and SIPEC (Marks et al., 2022) focus on behavior classification but do not provide quantitative measurements of the behavior intensity or the body kinematics during a behavior, which are key to behavioral analysis. Moreover, they are computationally expensive and require powerful graphics processing units (GPUs) to run.

Here, we report a new tool, *LabGym*, for quantifying user-defined animal behaviors without restrictions on behavior types or animal species, or the need to apply pre-defined or simplified features for behavior identification. When people manually analyze behaviors in a video, they typically ignore irrelevant information (such as static background) and make holistic observations of the behaving animal by considering both spatiotemporal changes and the overall movement pattern to categorize the behavior. Inspired by this human cognitive process, we designed *LabGym* to ignore static backgrounds, track multiple animals, accurately categorize behavior by holistic assessment, and then calculate quantitative measurements of each behavior. The technical innovations in *LabGym* include customizable combinations of neural networks for effective assessment of the spatiotemporal information in a behavior episode of user-defined duration, a new approach that we designed for efficiently evaluating the animal’s movement pattern during a behavior, and a new method for effective background subtraction. These innovations have made *LabGym* run easily on a common laptop computer, while most current deep-learning-based tools for behavioral analysis require demanding computational capabilities (Bohnslav *et al.*, 2021; Marks *et al.*, 2022; Mathis *et al.*, 2018).

Moreover, *LabGym* provides users a way to generate visualizable behavioral datasets that can be shared across the research community as ground truth (i.e., the data with assigned true labels) (LeCun et al., 2015; Roh et al., 2021; Schmidhuber, 2015) or for benchmarking different algorithms. Furthermore, *LabGym* includes real-time visualizations of behavioral categorizations and a graphical user interface (GUI) containing flexible options for experimenters to use without the need of writing computer code.

We show that *LabGym* precisely captures subtle changes in user-defined behaviors in animals ranging from soft-bodied invertebrates to mammals. We further demonstrate its strength by comparing it with the existing standards.

## RESULTS

### The *LabGym* pipeline

*LabGym* (https://github.com/umyelab/LabGym) was designed to mimic human assessment of animal behaviors. Our goals were to design software that efficiently and accurately identifies all the user-defined behavioral types (i.e., “categories”) for each animal in an experimental session, and that provides quantitative measurements of each category.

To accomplish this, *LabGym* first uses a new method to effectively remove background in each video frame and tracks each animal within the video (Figure 1A). At each frame of the video, a module termed the ‘Categorizer’ determines which behavior an animal performs during a time window that spans from the current frame back to a user-defined number of frames prior (‘time window’). To achieve this, two types of data are extracted at each video frame for each animal: a) a background-free animation (‘animation’) that spans the specified time window, and b) the imprint of the positional changes of the animal’s body over time in the animation (‘pattern image’; Figure 1B and Supplementary Video 1). These two pieces of data are then analyzed by the Categorizer that comprises three submodules: the ‘Animation Analyzer’, the ‘Pattern Recognizer’, and the ‘Decision Maker’ (Figure 1C). The Animation Analyzer analyzes all spatiotemporal details in each animation, whereas the Pattern Recognizer evaluates the animal’s overall motion pattern in the paired pattern image (see details in *“Biologically plausible design of the Categorizer for holistic assessments”*). The Decision Maker integrates the outputs from the Animation Analyzer and Pattern Recognizer to determine the behavior category within the specified time window.

**Figure 1.**
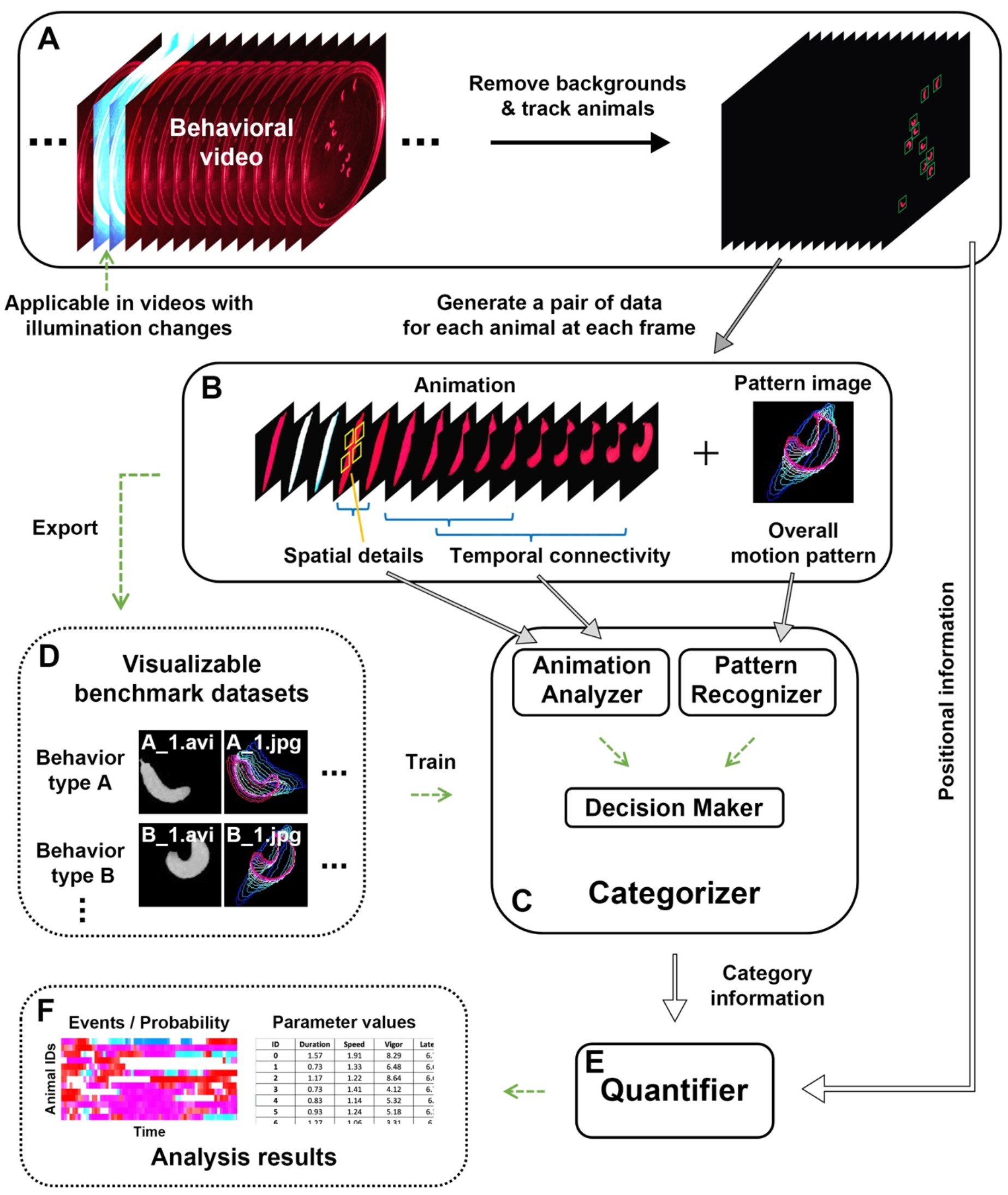
The pipeline of *LabGym*. **A**. Static backgrounds are removed in each video frame and individual animals are tracked. **B**. At each frame of the video, *LabGym* generates a pair of data for each animal, which comprises a short animation spanning a user-defined time window and its paired pattern image representing the movement pattern of the animal within the animation. **C**. The Categorizer uses both the animations and pattern images to categorize behaviors during the user-defined time window (i.e., the duration of the animation). The Categorizer comprises three submodules, the ‘Animation Analyzer’, the ‘Pattern Recognizer’, and the ‘Decision Maker’. The Animation Analyzer analyzes all spatiotemporal details in the animation for each animal whereas the Pattern Recognizer evaluates the overall movement patterns in the paired pattern image. The Decision Maker integrates the outputs from both the Animation Analyzer and the Pattern Recognizer to determine which user-defined behavioral category is present in the animation for each animal. **D**. The animations and pattern images can be exported to build visualizable, sharable benchmark datasets for training the Categorizer. **E** and **F**. After the behavioral categorizations, the quantification module (‘Quantifier’) uses information from the behavioral categories and the animal foregrounds to compute specific quantitative measurements for different aspects of each behavior.

Because it is based on deep neural networks, the Categorizer needs to be trained on examples of ground truth (i.e., user-labeled data) for each behavior in order to conduct accurate categorizations. To train the Categorizer, users only need to ‘tell’ the Categorizer what the behaviors are by providing examples (animations and their paired pattern images generated by *LabGym;* Figure 1B) of each behavior. To do so, users use *LabGym* to export unsorted example files, and manually sort each pair of example files into folders created under the behavior name (Figure 1D). Users then input these folders into *LabGym* to generate a labeled dataset for training the Categorizer.

After the categorizations, the quantification module called the ‘Quantifier’ (Figure 1E) uses information from the behavioral categories and the animal foregrounds to compute various parameters for each behavior (e.g., duration, speed, etc.; Figure 1F). The output also includes temporal raster plots of behavioral events (and their probabilities) and annotated videos for realtime visualization of behavioral categorization.

### Tracking multiple animals in diverse backgrounds for high-throughput and efficient analyses

Behavioral experiments often involve behavior-irrelevant backgrounds—such as the enclosures for holding the animals or changes in illumination during optogenetic manipulation—that renders analyses inaccurate and inefficient. To address these issues, we designed a module in *LabGym* that removes backgrounds but retains all the pixels representing the animals in each video frame (i.e., foreground) (See also below in *“Biologically plausible design of the Categorizer for holistic assessments”* for approaches for incorporating background information).

We reasoned that backgrounds in experimental conditions are often static, and that the pixel value in each background location would be stable over time (Supplementary Figure 1A). Therefore, we developed a method to find the stable value of each pixel in each video frame and use these values to reconstruct the static background (Figure 2A and see **Methods**). The static background is then subtracted from each frame to obtain the foreground, which represents the animal (Figure 2B). Compared with two state-of-the-art background subtraction methods, Local SVD Binary Pattern (LSBP) (Guo et al., 2016) and GSoC (Bradski, 2000) (implemented in Google Summer of Code 2017), our stable-value detection method reconstructed much cleaner backgrounds (with less traces of the animals) on behavioral videos of multiple animals and in diverse experimental settings (Figure 2B and Supplementary Figure 1B). Notably, animals are reliably tracked by *LabGym* even in videos with unstable or shifting illuminations (Supplementary Video 2) because the changes in background illumination (e.g., during optogenetic stimulation) can be reconstructed separately.

**Figure 2.**
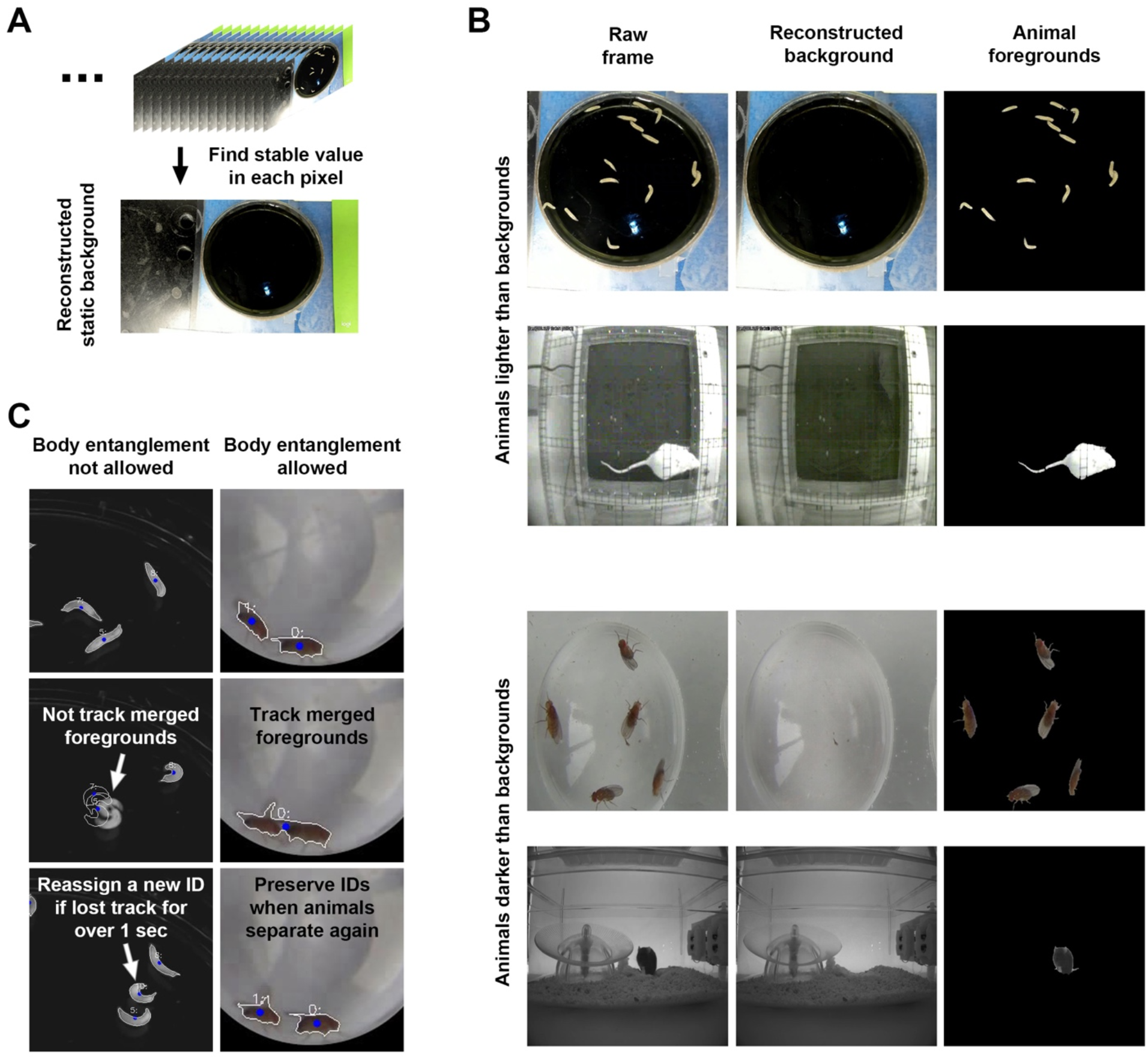
*LabGym* removes backgrounds and accurately tracks multiple animals. **A**. The stable value of each pixel is used to reconstruct the static background for a video. **B**. Examples of video frames showing that animal foregrounds are clearly presented after removing the static backgrounds that are constructed by our method of stable-value detection. **C**. Examples of video frames showing that *LabGym* accurately track multiple animals performing either non-interactive or social behavior. Left: non-interactive behavior, in which animal (larva) contact is not allowed; right: social interaction in which animal (adult fly) contact is allowed.

The tracking of each animal over time is performed based on the assumption that the Euclidean distance between the centers of the same animal in consecutive frames are always the smallest among all Euclidean distances between each possible pair of animal centers. Each tracked animal is assigned by a unique identity (ID) that links to a matrix for storing all information about that animal over time.

Multiple animals in the same enclosure sometimes collide with each other, which can cause tracking errors. To make the analysis fully automatic, the current version of *LabGym* excludes animals from tracking if users choose not to allow animal body entanglements (Figure 2C, Supplementary Videos 3 and **Methods**). This ensures that the analysis is only performed on reliably tracked individual animals. If body entanglements are allowed, *LabGym* can also analyze social behaviors in which animals have body contact (Figure 2C, Supplementary Videos 4 and **Methods**). In this scenario, merged animals are analyzed as one.

Taken together, the effective background subtraction and reliable animal tracking in *LabGym* make it applicable to a wide range of behavioral videos without the need for specialized recording equipment.

### Biologically plausible design of the Categorizer for holistic assessments

We built the Categorizer with three submodules: Animation Analyzer, Pattern Recognizer, and Decision Maker (Figure 3 and **Methods**), each of which consists of neural networks that suit the design goals. Importantly, the two subnetworks—the Animation Analyzer and the Pattern Recognizer—were designed to analyze two types of behavioral information that complement each other, which ensures accurate categorizations of behavior through holistic assessment.

**Figure 3.**
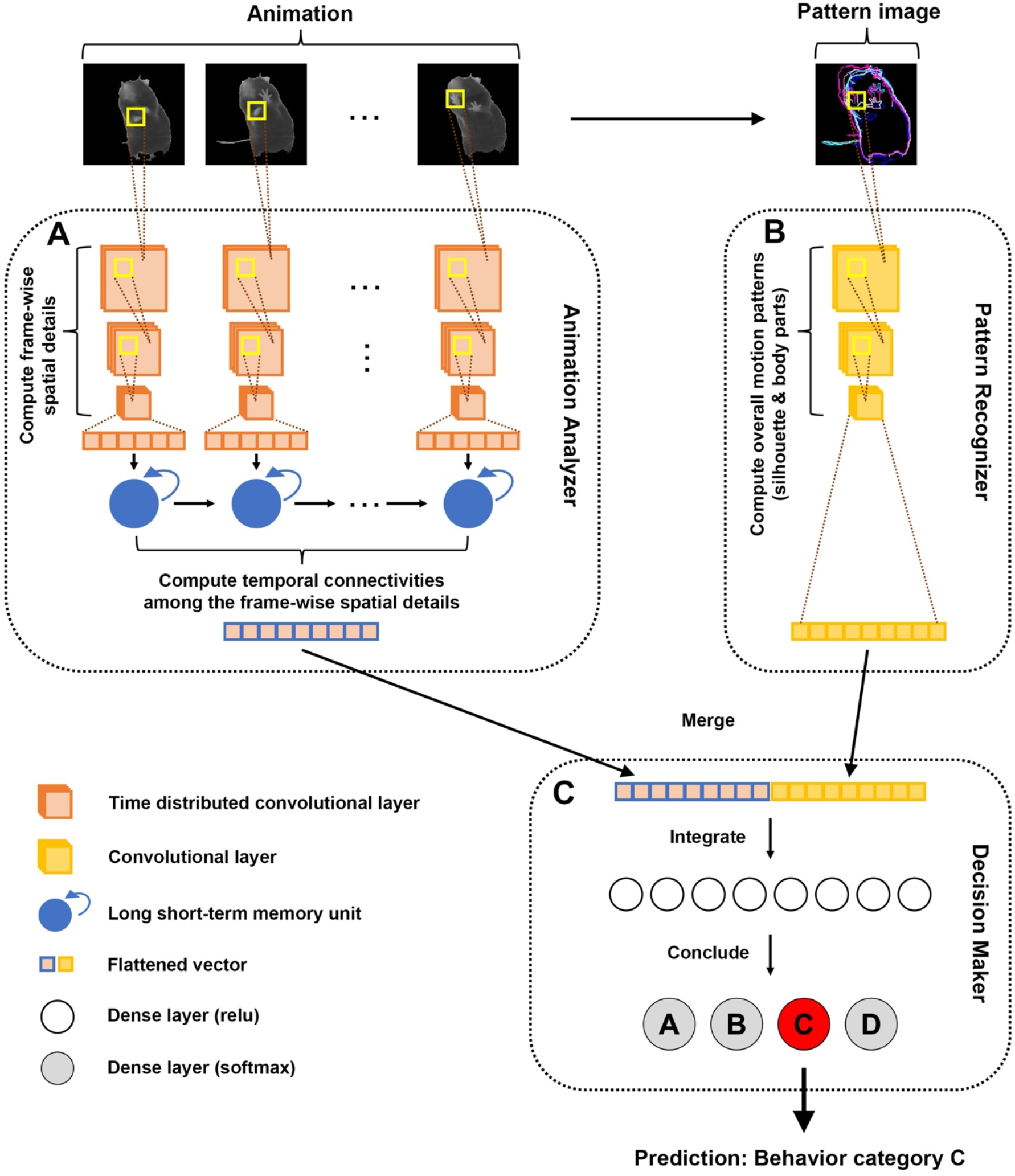
Biological plausible design of the Categorizer. **A**. The Animation Analyzer first uses time-distributed convolutional layers to compute all frame-wise spatial details of an animation, and then uses recurrent layers (long short-term memory, LSTM) to compute the temporal connectivity among the frame-wise spatial details. **B**. The Pattern Recognizer uses convolutional layers to analyze the pattern image, which is superimposed contours of animal body (and body parts) along the temporal sequence (indicated by gradually changed color) during a behavior. **C**. The Decision Maker uses concatenating layers to integrate the outputs from both the Animation Analyzer and the Pattern Recognizer, and then passes the integrated information to dense layers for concluding the behavioral categories.

Animal behavior contains changes in both spatial and temporal dimensions. To learn the spatial details of the animal in each time point (frame) during a behavior, the Animation Analyzer uses time-distributed convolutional layers to analyze raw pixel values that represent the animal foreground in each frame of an animation (Figure 3A). To learn about how these frame-wise spatial details are organized along the temporal axis, the Animation Analyzer then uses recurrent layers to compute temporal connectivity among the frame-wise spatial details analyzed by time-distributed convolutional layers.

When an animal performs a behavior, the positional changes of its body and body parts over time can form a pattern that is unique to the behavior. This pattern is key to behavioral categorizations. To enable the Categorizer to capture distinct patterns, a “pattern image” was generated for each animation to depict the animal’s overall movement and the Pattern Recognizer (that comprises convolutional layers) was designed to analyze the pattern image (Figure 3B). In each pattern image, the contours of both the whole body and the body parts of the animal in each frame of the animation were superimposed with gradually changed colors to indicate their temporal sequences. This shows the animal’s overall movement pattern during the animation in a single 2-dimensional image to allow for efficient analyses.

Finally, the Decision Maker uses concatenating layers to integrate the outputs from both the Animation Analyzer and the Pattern Recognizer, and passes the integrated information to fully-connected (dense) layers for discriminating behavioral categories (Figure 3C).

Users can choose to include background in the animations as this sometimes can be useful for behavioral categorization. For example, bedding in a mouse’s mouth can indicate nest building and, thus, including the bedding in the animations can facilitate the categorization of nest building behavior (Figure 4A). Users can also choose whether to show the animals’ body parts in the pattern images. Whole-body motion patterns are sometimes sufficient for distinguishing behaviors like curling versus uncoiling in *Drosophila* larvae, wing extension versus abdomen bending in adult flies, or walking versus rearing in rats (Figure 4B), while movement patterns of individual body parts are key for differentiating similar behaviors like facial grooming versus sniffing, or ear grooming versus sitting still in mice (Figure 4C).

**Figure 4.**
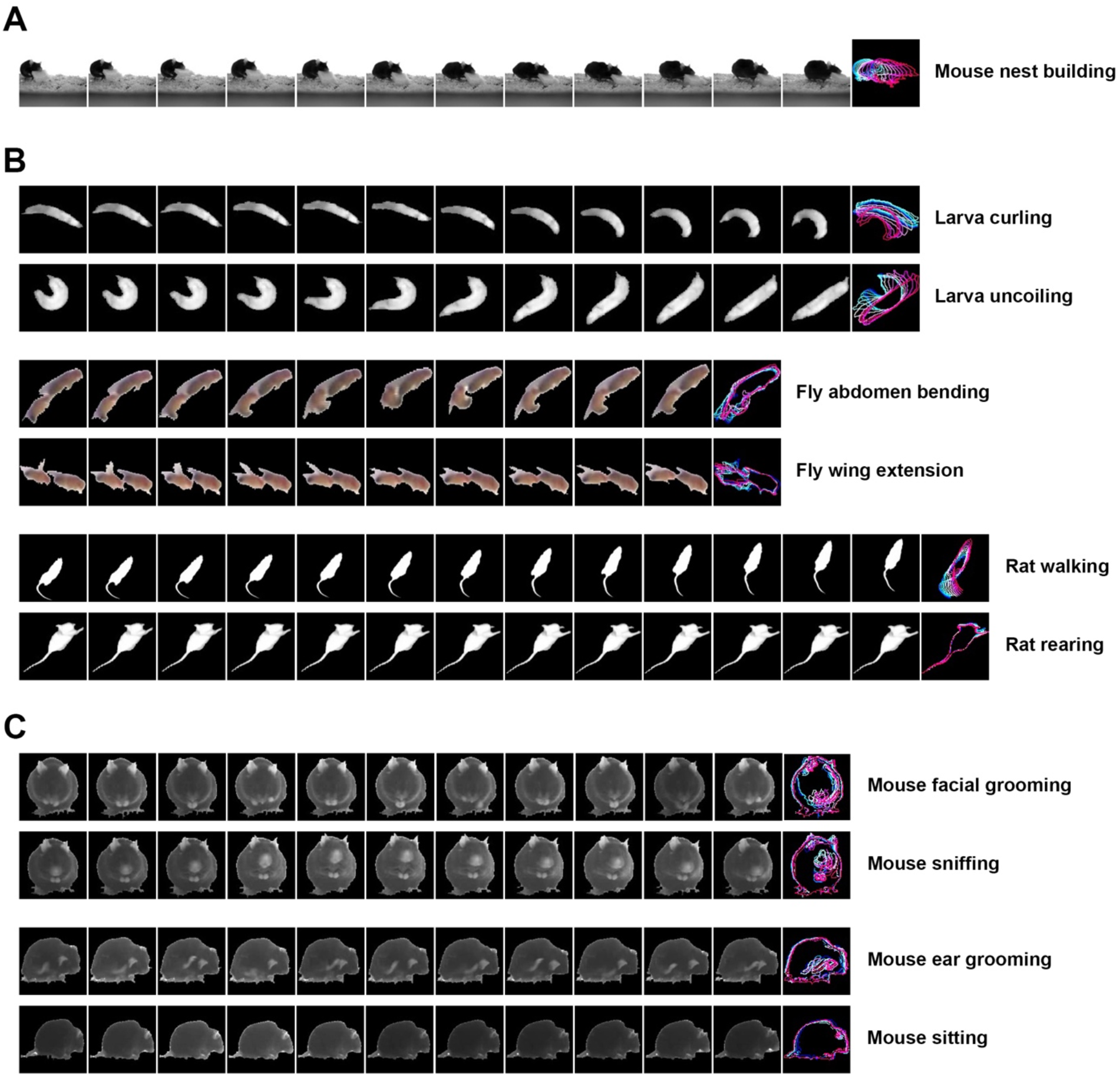
*LabGym* generates stand-alone, visualizable behavioral benchmark datasets. **A**. Example frames of an animation (background is included) and its paired pattern image, showing nest building behavior of a mouse. **B**. Example frames of animations and their paired pattern images (body parts are not included), showing (from top to bottom) larva curling and uncoiling, fly abdomen bending (copulation attempt) and wing extension (courtship song), and rat walking and rearing. **C**. Example frames of animations and their paired pattern images (body parts are included), showing (from top to bottom) mouse facial grooming, sniffing, ear grooming, and sitting.

### The Categorizer accurately categorizes user-defined behaviors across species

Since the behavior types for the Categorizer to recognize are defined by users, rather than pre-defined in the program, the Categorizer can identify behaviors without restrictions on behavioral categories or animal species. To demonstrate this, we used *LabGym* to study relatively simple behaviors in *Drosophila* larvae and behaviors of increasing complexity in rodents. We first used *LabGym* to generate and label separate behavioral datasets for *Drosophila* larvae (Supplementary Video 5), rats (Supplementary Video 6), and mice (Supplementary Video 7). These datasets are publicly available (see **Methods**), can be used by others, and can be modified and expanded on by other users to suit their studies. The larva dataset contains the fewest categories with sufficient examples to cover almost all the variations of each behavior; the rat dataset contains a larger number of categories and more complex behaviors that encompass subtle distinctions between categories; the mouse dataset contains additional categories of even more complex behaviors. To test the robustness of *LabGym*, we also varied the amount of training examples used to categorize a given behavior, ranging from ~100 to ~800 pairs of animations and pattern images.

The Categorizer is customizable with different levels of complexity to address behaviors of various complexity. To provide a reference for customizing the Categorizer, which is a trial- and-error process, we trained several Categorizers with different complexity for each species, and determined the most suitable ones based on their overall accuracy of behavioral categorizations in training (Supplementary Table 1 and **Methods**). The selected Categorizer for each dataset was named as *LarvaNociceptor_Topview_30fps* (*LarvaN*), *RatAddiction_Topview_30fps* (*RatA*) and *MouseHomecage_Sideview_30fps* (*MouseH*). To test the accuracy and generalizability of these three Categorizers, we assessed their precisions, recalls (sensitivity), and F1 score (weighted precision and sensitivity, see **Methods**) for behavioral categorizations of videos that were not used for generating the training or validation dataset.

Complex deep neural networks might not be suitable for simple larval behaviors since they would learn too many irrelevant details in simple datasets, resulting in reduced accuracy (i.e., overfitting). In fact, *LarvaN* with simple complexity (Supplementary Table 1) achieved 97% accuracy in frame-wise categorizations of 6 different behaviors elicited by noxious stimuli (Figure 5A and Supplementary Video 8).

**Figure 5.**
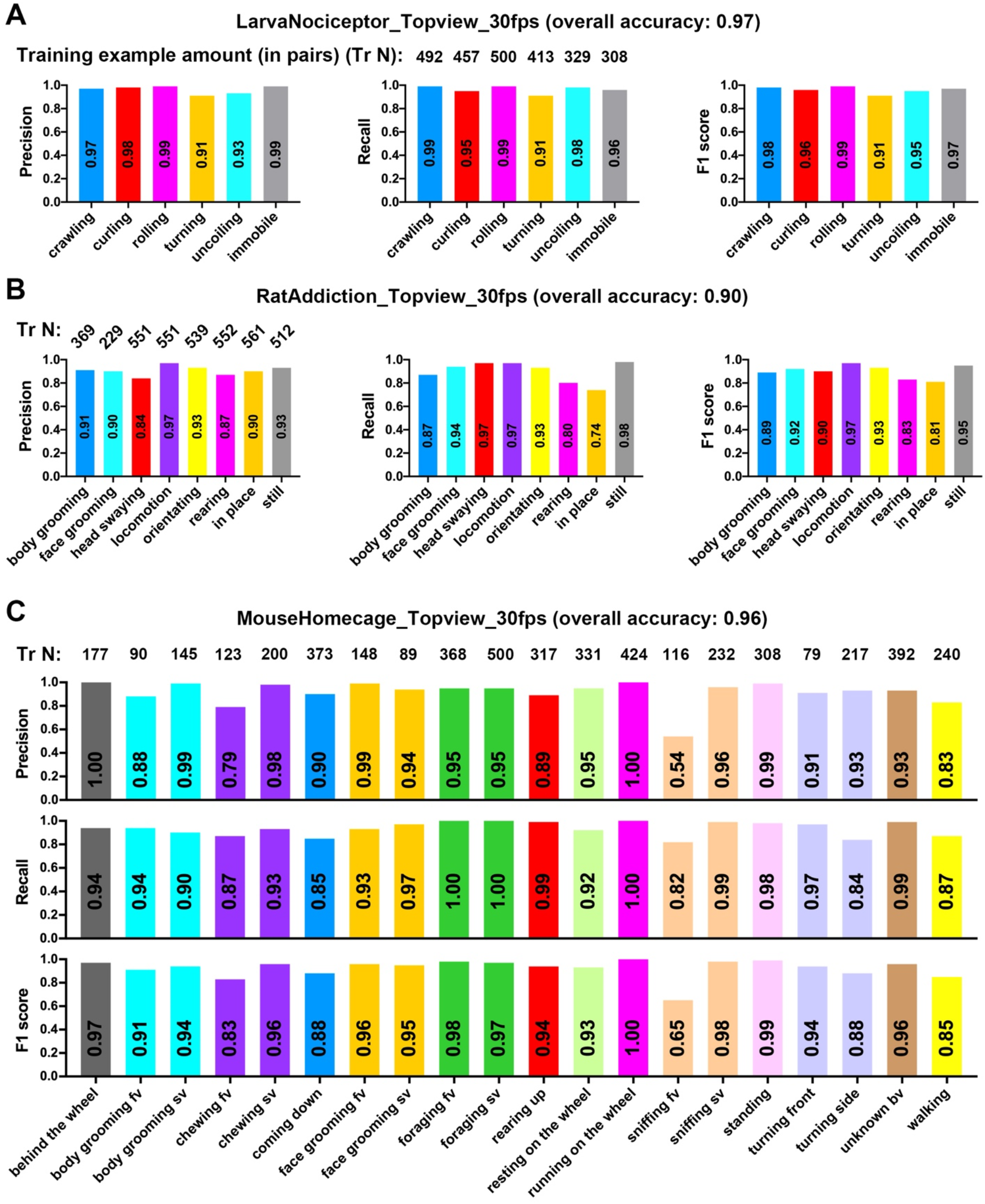
Accurate frame-wise categorizations of various user-defined behaviors by different Categorizers. **A-C**. Bars showing the in-practice test metrics (overall accuracy, precisions, recalls and F1 scores) for *LarvaN* (**A**), *RatA* (**B**) and *MouseH* (**C**) in video analysis. 7 videos for larvae (10 larvae per video), 10 videos for rats (1 rat per video), and 2 videos for mice (1 mouse per video) that were not used for generating the training datasets were used to generate the testing data (animations and their paired pattern images). 2162 pairs (for larvae), 4828 pairs (rats), and 12653 pairs (mice) of data were randomly selected (data with visibly missing body parts were excluded) and then were sorted by experimenters into different folders under user-defined behavioral names (building ground truth testing datasets). The ground truth testing datasets were then used to test the categorization metrics of the 3 Categorizers. In this way, experimenters were blind to the predictions of the 3 Categorizers when sorting data to build the ground truth testing datasets and thus the tests were unbiased.

Rats show complex behaviors and the differences among them can be extremely subtle due to the recording or experimental conditions (Supplementary Video 6). Despite these challenges, the *RatA* performed reliably well in frame-wise differentiation of visually similar behaviors such as body grooming versus face grooming, or head swaying versus in place (Figure 5B and Supplementary Video 9).

Animals can develop new behaviors when experimental conditions change. For example, mice in a cage equipped with a wheel tend to run on the wheel. The movement of running on the wheel has features and kinematics that are distinct from running across a flat surface. Thus, wheel-running might not be correctly identified by tools using predefined, generalized definition of running. Additionally, a camera placed at the side of a cage will result in at least two viewpoints of the mouse at it moves in the cage, a side view and a front view, which can greatly complicate the automated analysis of behavior. Despite these challenges, *MouseH* achieved excellent accuracy in frame-wise categorization of 20 behavioral categories (especially for wheel-running, which achieved 100% precision and recall) with as few as 79 training examples for some categories (Figure 5C and Supplementary Video 10). Although both were used for categorizing rodent behaviors, *MouseH* trained on pattern images including the body parts (Supplementary Videos 7) performed better than *RatA* trained on those excluding body parts (Supplementary Videos 6; Figure 5B and 5C). This indicates that including the body parts in pattern images can boost the learning efficiency of the Categorizer (Figure 4 and **Methods**).

Due to the data-driven nature of deep learning, the accuracy and generalizability of the Categorizer largely depends on the amount, diversity, and labeling accuracy of the training examples. Therefore, the performance of the Categorizers can be continuously improved over time with the accumulation of well-labeled training examples.

### The Quantifier provides quantitative measurements that are sufficient for capturing subtle changes in diverse aspects of each behavior

To quantitatively measure the behavior intensity and the body kinematics during a behavior, we designed a quantification module in *LabGym* called ‘Quantifier’. The Quantifier calculates 14 parameters to quantify the behavior: count, latency, duration, angle, speed (distance traveled per time unit), velocity (displacement per time unit), acceleration/velocity reduction, distance, intensity (area), intensity (length), magnitude (area), magnitude (length), vigor (area) and vigor (length) (Figure 6 and **Methods**). Additional parameters can be added in future versions of *LabGym*. The definitions of these parameters are:

- The **count** is the summary of the behavioral frequencies, which is the occurrence number of a behavior within the entire duration of analysis. Consecutive single occurrences (at a single frame) of the same behavior are considered as one count.
- The **latency** is the summary of how soon a behavior starts, which is the time starting from the beginning of the analysis to the time point that the behavior occurs for the first time.
- The **duration** is the summary of how persistent a behavior is, which is the total time of a behavior within the entire duration of analysis.
- The **angle** is the movement direction (against to the animal body axis) of the animal during a behavior episode, which is the mean of all the included angle (*θ*) between animal body axis and the movement direction during the time window (*t_w_*) for categorizing the behavior.
- The **speed** is the summary of how fast the animal moves when performing a behavior, which is the total distance traveled (*d*) (between the two centers of mass of the animal) during the time window (*t_w_*) for categorizing the behavior divided by *t_w_*.
- The **velocity** is the summary of how efficient the animal’s movement is when performing a behavior, which is the maximum displacement (*dt*) (between the two centers of mass of the animal) divided by the time (*t*) that such displacement takes place.
- The **acceleration** / **velocity reduction** is the summary of how fast the animal’s velocity changes while performing a behavior, which is the difference between maximum velocity (*v_max_*) and minimum velocity (*v_min_*) divided by the time (*t_v_*) that such velocity change takes place.
- The **distance** is the total distance traveled of the animal by performing a behavior within the entire duration of analysis.
- The **intensity (area)** / **intensity (length)** is the summary of how intense a behavior is, which is the accumulated proportional changes of the animal body area (*a*) / length (*l*) between frames divided by the time window for categorizing the behaviors (*t_w_*) when performing a behavior.
- The **magnitude (area)** / **magnitude (length)** is the summary of the motion magnitude, which is the maximum proportional change in animal body area (*a*) or length (*l*) when performing a behavior.
- The **vigor (area)** / **vigor (length)** is the summary of how vigorous a behavior is, which is the magnitude (area) / magnitude (length) divided by the time (*t_a_* or *t_l_*) that such a change takes place.

**Figure 6.**
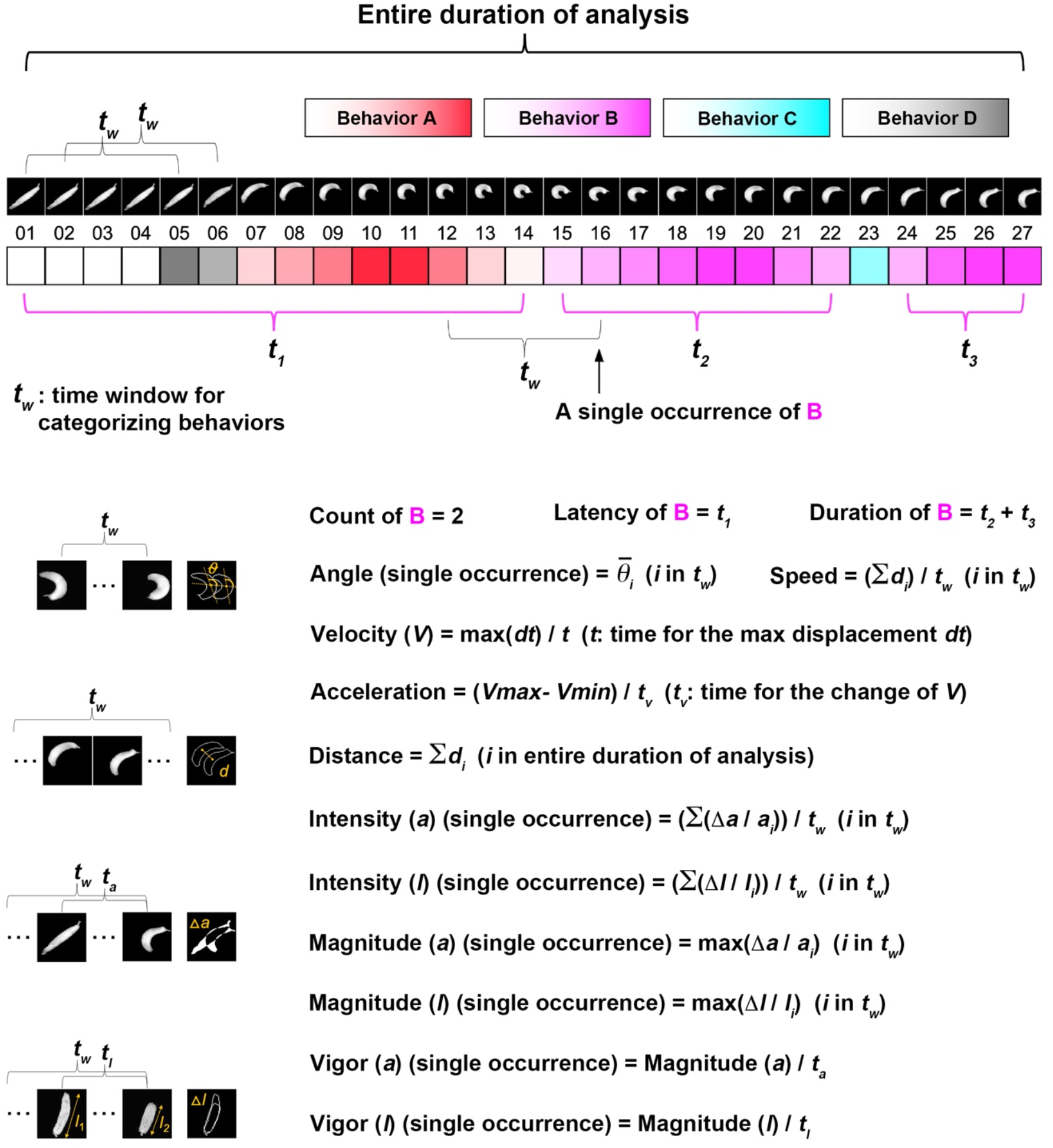
The Quantifier analyzes diverse aspects of a behavior. A schematic that shows the process of frame-wise categorizations on behaviors, and how the Quantifier uses the information of behavioral categories and animal foregrounds to calculate 14 behavioral parameters in *LabGym*.

To test how well the Quantifier performs in quantifying behaviors in diverse animal species, we applied *LabGym* to experiments in larvae, rats, and mice, in which the quantitative measurements of behaviors was difficult to achieve with previous methods.

#### Drosophila larva behavior

Previous studies show that optogenetic inhibition of larval leucokinin (LK) neurons slightly impaired larval behaviors elicited by nociceptor activation (Hu et al., 2020). In these experiments, both the subtle changes in behaviors and the shifting illumination caused by optogenetic manipulations posed challenges for automatic behavioral analysis. We thus tested whether *LabGym* was able to capture the behavioral changes in these experiments (Supplementary Video 11). Compared with the control group, larvae in the LK inhibition group showed a modest reduction in the probability of body rolling behavior, which is a characteristic nociceptive response (Hwang et al., 2007; Tracey et al., 2003) (Figure 7A and 7B). No change was observed in the latency of the response onset after optogenetic stimulation of nociceptors (Figure 7C). Moreover, LK inhibition caused a slight yet significant reduction in rolling velocity (Figure 7D) and duration (Figure 7E). We also compared several quantitative measurements of different behaviors and found that the results indeed reflect the ethological meaning of these behaviors. For example, rolling achieves higher speed than crawling (Figure 7F), supporting the notion that rolling was a more efficient way than crawling to escape from harm (Hwang *et al.*, 2007; Tracey *et al.*, 2003). Similarly, despite their spatial similarity, curling was more vigorous than turning (Figure 7G), echoing the fact that larvae typically perform curling under noxious stimuli but tend to turn in in response to milder situation like gentle touch or exploration (Hwang *et al.*, 2007; Jovanic et al., 2016; Lahiri et al., 2011; Tracey *et al.*, 2003). Thus, *LabGym* is sufficiently sensitive to detect modest effects of in vivo neuronal inhibition on behavior, and it does so on videos with shifting illumination.

**Figure 7.**
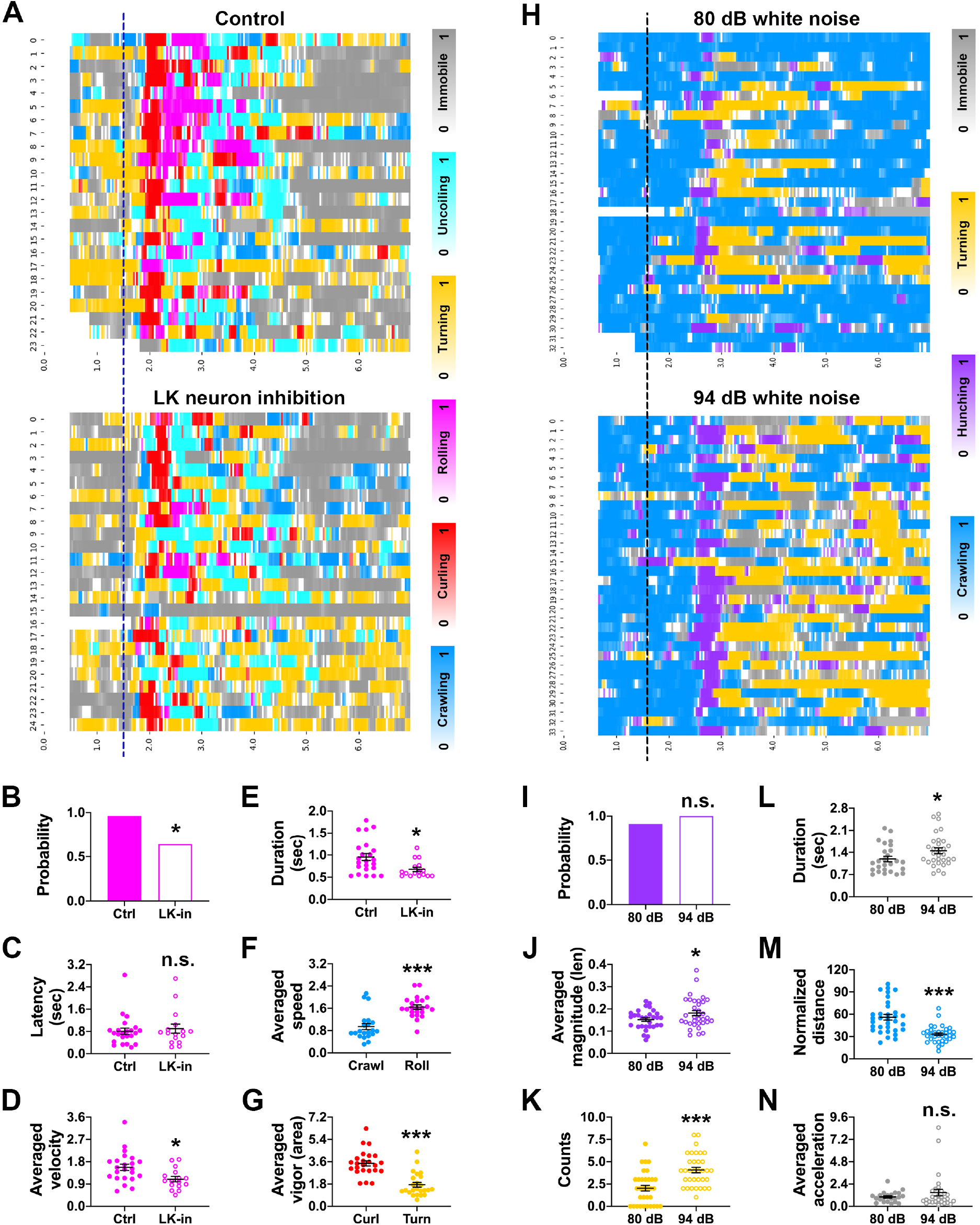
The Quantifier captures subtle changes in the behavior of *Drosophila* larvae. **A** and **H**. Raster plots showing the frame-wise categorizations by: **A**. *LarvaN* on control (*w*; *LK*-GAL4 / +; *TrpA1*-QF, QUAS-ChR2^T159C^, UAS-jRCaMP1b / +) and LK neuron inhibition (*w*; *LK*-GAL4 / +; *TrpA1*-QF, QUAS-ChR2^T159C^, UAS-GtACR1 / +) groups after nociceptors were optogenetically activated (blue dashed line indicates the stimulation onset); **H**. *LarvaMechanosensor_Topview_30fps* on larvae (*Caton S*) under different intensities of sound stimuli (black dashed line indicates the stimulation onset). Color bars: behavioral categories (color intensities indicate the behavioral probabilities). X-axis represents the time and Y-axis represents the larval IDs. **B-G**. Parameters calculated by the Quantifier for different behaviors elicited by optogenetic activation of nociceptors. The colors match those for behaviors shown in **A**. Data in **F** and **G** are from the control group in **A**. **I-N**. Parameters calculated by the Quantifier for different behaviors elicited by sound stimulation. The colors match those in **H**. Probability in **B** and **I** is the fraction of responders (behavioral count > 0) in total animals. n.s., P > 0.05; *, P < 0.05; ***, P < 0.001 (Fisher’s exact test for **B** and **I**; unpaired t test for **C-G** and **J-N**).

When *Drosophila* larvae are exposed to sound, they display a startle response that is characterized by the cessation of crawling (“freezing”), retraction of their heads (“hunching”), and excessive turning movements (Zhang et al., 2013). To test whether *LabGym* could identify the subtle differences in behaviors elicited by different intensities of sound, we trained another Categorizer *(LarvaMechanosensor_TopView_30fps)* with examples of sound-elicited behaviors in *Drosophila* larvae (Supplementary Videos 5) and used it for analysis (Supplementary Video 12). This Categorizer has the same complexity as *LarvaN*. Compared to larvae exposed to 80 decibel (dB) white noise, those exposed to 94 dB showed a similar hunching probability (Figure 7H and 7I), but the magnitude of head retraction during hunching was enhanced (Figure 7I). Furthermore, larvae exposed to 94 dB showed more frequent turning (Figure 7K) and prolonged freezing (i.e., immobility; Figure 7I), which resulted in significant reductions in crawling distance (Figure 7M). However, when larvae froze, the reduction in the velocity of their movement was similar between the two groups (Figure 7N). Thus, *LabGym* was able to capture quantitative differences in the behaviors induced by auditory stimuli of different intensities.

#### Rodent behavior

It is well-established that drugs like amphetamine, produce dose-dependent increases in psychomotor activity, and that the intensity of these behaviors increases (i.e., sensitizes) with repeated drug exposure (Ellinwood and Balster, 1974; Ferrario *et al.*, 2005). Psychomotor activity in rats encompasses a wide range of behaviors that include rearing, hyper-locomotion, and stereotyped, repetitive head movements (i.e., stereotypy). Psychomotor activity is most often quantified using non-parametric rating scales based on clusters of behavioral features and their intensities (Ellinwood and Balster, 1974; Ferrario *et al.*, 2005). To date, no automated systems have been able to sufficiently categorize or quantify psychomotor activity, despite early efforts that predate machine learning (Flagel and Robinson, 2007).

*LabGym* automatically captured dose-dependent changes in key behavioral features of psychomotor activity (Figure 8A and Supplementary Video 13). These include rearing (Figure 8B), locomotion (Figure 8C), and head movements (persistent head movements indicating stereotypy; Figure 8D), following systemic amphetamine administration. Impressively, *LabGym* even captured stereotypy interspersed with small bursts of locomotion—an indication of a strong psychomotor response at high dose of amphetamine (e.g., Figure 8A and 8C, 5.6 mg/kg dose group in amphetamine baseline group). *LabGym* also captured quantitative changes in these behaviors across the development of sensitization, in which repeated injection of amphetamine resulted in more intense behavioral responses to the same dose of drug (Figure 8E–8H). In contrast, the intensity of spontaneously occurring behaviors such as body/face grooming were stable across time in rats treated with repeated saline (Figure 8I–8K).

**Figure 8.**
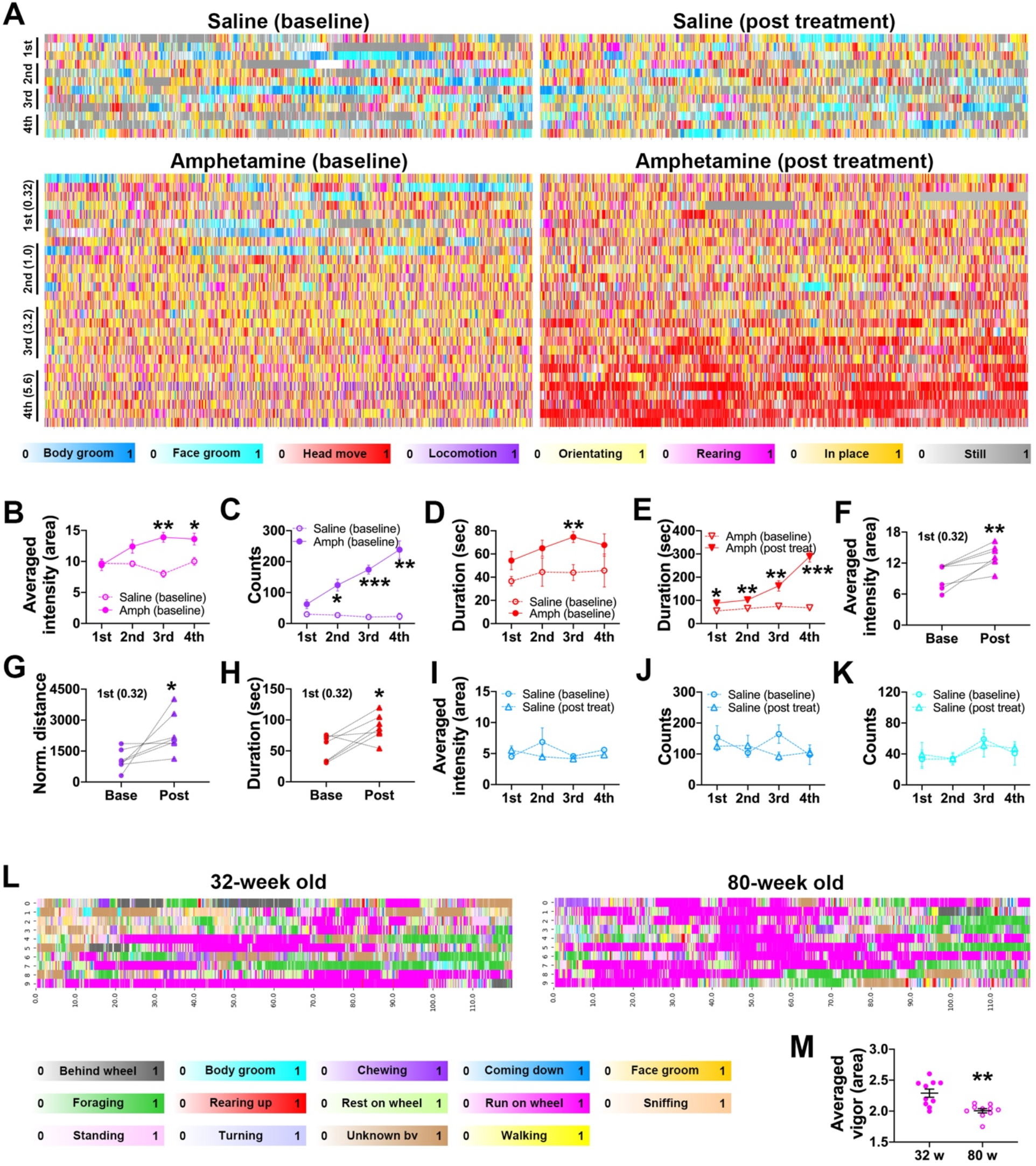
The Quantifier captures qualitative and quantitative changes in rodent behaviors. **A**. Raster plots showing the frame-wise categorizations by *RatA* of behavior after saline or amphetamine injection. Plots at the left (baseline) show responses to the first exposure to amphetamine across increasing doses (baseline; 0.32, 1.0, 3.2, and 5.6 mg / kg, i.p.); Plots at the right show responses to increasing doses of amphetamine after repeated amphetamine treatment (post treatment; 21 days, see methods). Animals in the saline group received repeated saline throughout. Color bars: behavioral categories (intensities indicate the behavioral probabilities). X-axis: time (each tick is 10-sec); Y-axis: each row represents an individual rat. These same colors are used to indicate behavioral categories in panels **B-K**. **B-D**. The dose response curves of rearing intensity (**B**), locomotion counts (**C**) and head movement duration (**D**) following the first exposure to amphetamine or saline (baseline). *, P < 0.05; **, P < 0.01; ***, P < 0.001 (unpaired t test between saline and amphetamine groups). **E**. Dose response curves for head movement duration at baseline and after repeated amphetamine exposure (post treatment, i.e., sensitization). *, P < 0.05; **, P < 0.01; ***, P < 0.001 (unpaired t test between baseline and post treatment groups). **F-H**. Within subject comparison of rearing intensity (**F**), locomotion distance (**G**) and the head movement duration (**H**) in response to the first 0.32 mg / kg amphetamine at baseline and after repeated amphetamine treatment. *, P < 0.05 (paired t test). **I-K**. Dose response curves of the intensity and counts of body grooming (**I** and **J**) and the duration of face grooming (**K**) in in rats repeatedly treated with saline. **L**. Raster plots showing the frame-wise categorizations by *MouseH* for a younger (32-week) or an older (80-week) mouse. Color bars: behavioral categories (intensities indicate the behavioral probabilities). X-axis: time (in second); Y-axis: different random 2-minute time window. Multiple (>100) 2-minute time windows were randomly selected from a 4-hour recording for each mouse, and those containing wheel-running behavior (targeted time window) were sorted out. 10 targeted time windows for each mouse were randomly selected for analysis. **M**. The running vigor of the two mice in **L**. **, P < 0.01 (unpaired t test).

When an animal ages, its ethogram might not change, yet the quantitative measurements of its behaviors (e.g., running speed) may decline. The speed of a mouse running on a wheel is typically measured by tracking the rotation of the wheel itself, which poses a challenge for computational tools designed to only track animals. Nevertheless, *LabGym* not only categorized the all the behaviors of interest for both younger (32-week) and older mice (80-week) in a cage with a running wheel (Figure 8L and Supplementary Video 14), but also captured the decline in movement speed of the older versus the young mice by a parameter (vigor_area) that summarizes the areal changes per unit time of the animal’s body (Figure 8M, 6 and **Methods**).

Taken together, these results demonstrate that the Quantifier reliably and precisely captures quantitative changes in behaviors across species, and in a wide range of settings and experimental conditions.

### GUI with optional settings for user-friendly usage

To make *LabGym* adaptable to various experimental settings and accessible to users without requiring a computer programming background, we implemented a GUI. This includes 4 functional units: Generate Datasets, Train Networks, Test Networks and Analyze Behaviors.

The Generate Datasets unit (Supplementary Figure 2) is for batch processing of videos to generate visualizable behavioral data (animations and paired pattern images) that can be used to establish benchmark datasets for training the neural networks. The Train Networks unit (Supplementary Figure 3) is for preparing training datasets and training Categorizers at various levels of complexity. The Test Networks unit (Supplementary Figure 4) is for testing the accuracy and generalizability of a trained Categorizer in an unbiased manner. The Analyze Behaviors unit (Supplementary Figure 5) is for batch-processing of videos for behavioral analysis. It generates an annotated video (Supplementary Videos 8-14), a raster plot of behavioral events (displayed in user-defined colors) and their occurring probabilities (by color intensities) for each animal in one batch of analysis (Figure 7A, 7H, 8A and 8L), and a spreadsheet in Excel or CSV format containing individual behavioral measures for each behavior of each animal.

Notably, the flexible options in *LabGym*, especially the customizable neural network complexity, allows it to achieve fast processing speed in training networks and in analyzing behaviors on commonly used laptops that do not have GPUs (Supplementary Table 2). This is difficult to achieve for tools using fixed and complex neural networks (Bohnslav *et al.*, 2021; Mathis *et al.*, 2018).

### Benchmark comparison with the existing standards

The combination of deep learning-based tracking tools and the post-hoc behavior analysis tools can achieve multi-animal tracking, classification and quantification of user-defined behaviors in different animal species, while *LabGym* can achieve the same goal in one tool. We directly benchmarked the performance of *LabGym* against that of the combination of DeepLabCut (DLC) (Mathis *et al.*, 2018) and B-SOiD (Hsu and Yttri, 2021) (“DLC+B-SOiD”), which represent the existing standards and also have GUIs for accessibility to users without requiring computer programming knowledge.

All the three tools can be iteratively optimized by adding more training data and refining the data labeling. Therefore, the goal of the benchmark comparison is not to explore how to modify the training datasets to achieve the best performance of these tools, but to compare their performance with the same training resources and labor workload (using a fixed 30-minute for manual labeling and sorting for both *LabGym* and DLC+B-SOiD). To achieve this, we used videos in which animals display behaviors of interest that differ in intensity. We then determined whether each approach (DLC+B-SOiD and *LabGym)* can detect the behaviors of interest, and how results compare across these approaches (Figure 9A). Specifically, the benchmarking training dataset used contained 4 videos (2-minute duration at 30 fps) covering major types of rat behavior in psychomotor activity experiments: locomotion, rearing, repeated head movements and other background behaviors such as resting (in place) and orientating. The labor for DLC+B-SOiD was spent on labeling animal body parts in extracted frames for tracking (~25 minutes) and assigning behavior names for behavior classification (~5 minutes), while that in *LabGym* was spent on sorting behavior data for behavior classification (~30 minutes). The testing dataset contained 3 videos (1-minute duration at 30 fps per video):

1. A video (referred as ‘Base’) that was randomly selected and trimmed from ‘Amphetamine (baseline)’ in Figure 8A, in which the rats were exposed to 5.6 mg/kg amphetamine for the first time. The rat in this video showed psychomotor activities such as frequent rearing and locomotion.
2. A video (referred as ‘Post’) that was randomly selected and trimmed from ‘Amphetamine (post treatment)’ in Figure 8A, in which the rats were exposed to 5.6 mg/kg amphetamine after experiencing repeated amphetamine exposure (i.e., a sensitizing regimen). The rat in this video showed intensified psychomotor activity such as frequent stereotypy (repeated head moving).
3. A video from (Ferrario *et al.*, 2005) in which testing occurred in a different enclosure than that used in the training videos. This video was used to test the generalizability of these tools.

**Figure 9.**
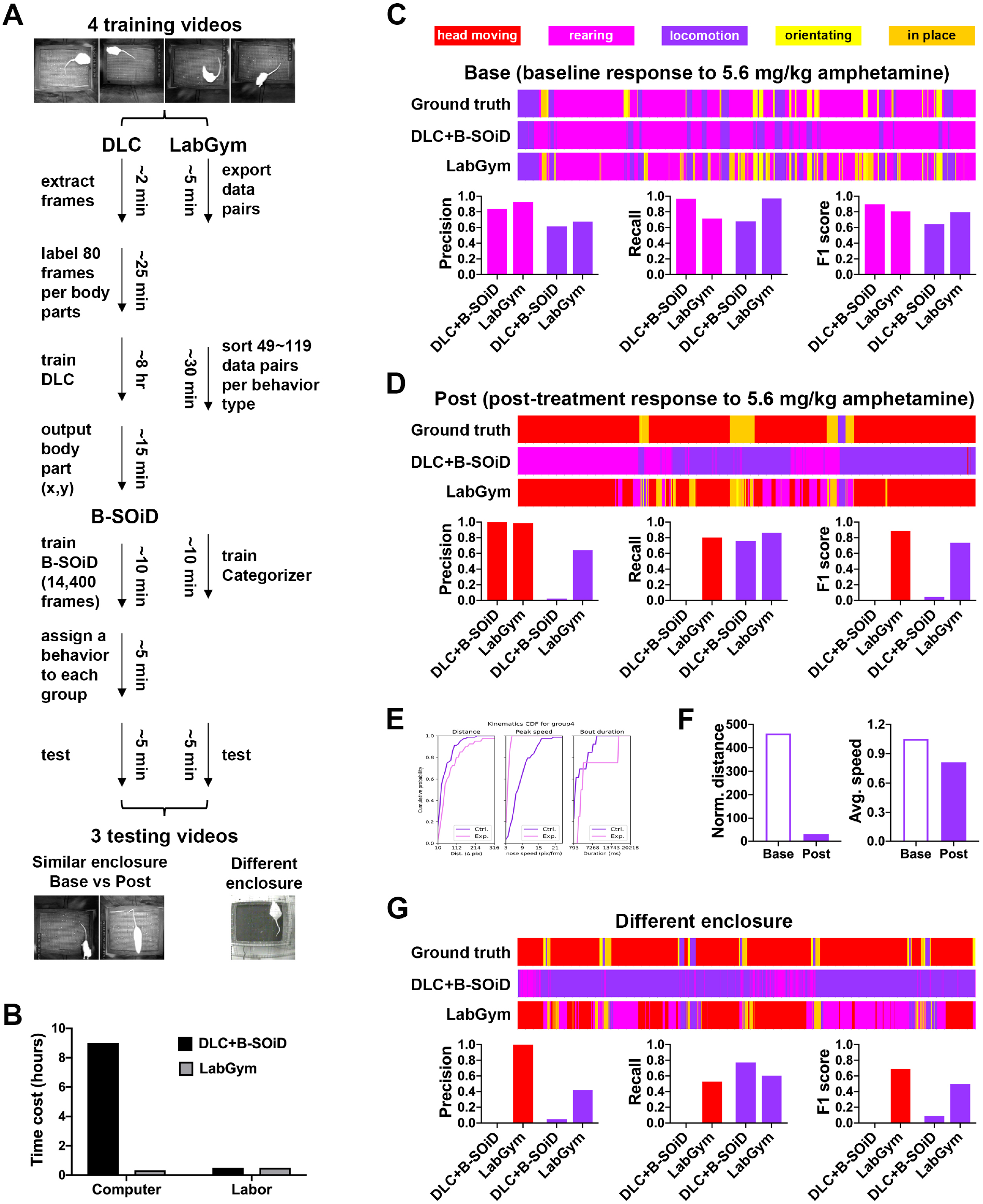
Benchmark comparison between the existing standards and *LabGym*. **A**. The workflow of the benchmark comparison between the combination of DeepLabCut and B-SOiD (DLC+B-SOiD) and *LabGym*, which was performed on the same computer that has a 3.6 GHz Intel Core i9-9900K CPU, a 64 GB memory, and an NVIDIA GeForce RTX 2080 GPU). The setting of Categorizer in *LabGym:* Animation Analyzer: level 1 with input shape 8 X 8 X 1; Pattern Recognizer: level 2 with input shape 16 X 16 X 3. **B**. The total time spent for DLC+B-SOiD and *LabGym* to finish the workflow in **A**. The total human labor was fixed to 30 minutes for both. **C-D**. Comparisons of frame-wise behavior classifications between the computer software and the ground truth (expert human annotation) in 2 testing videos: baseline (**C**) and post-treatment (**D**). The raster plots show frame-wise behavior events with the X-axis indicating the time (1 minute). Different colors indicate different behavior types. Videos used here were randomly selected from those used for Figures 8**A**-8**E**. **E-F**. Examples of behavior quantification (locomotion) in baseline (base) and post-treatment (post) videos by B-SOiD (Ctrl = baseline, Exp = post-treatment; **F**) and *LabGym* (**G**). **G**. Comparison of frame-wise behavior classifications between the computer software and the ground truth (expert human annotation) in generalization to a different testing enclosure. The raster plots show frame-wise behavior events with the X-axis indicating the time (1 minute). The colors match those for behaviors shown in **C**.

Comparisons were performed on two key aspects: the total time cost and the end accuracy of identifying and quantifying behaviors of interest. We first input the 4 training videos (8 minutes in total at 30 fps) into DLC to extract frames (~2 minutes) and labeled 6 body parts in each frame (~25 minutes). We then used these labeled frames to train a DLC model (~8 hours). This resulted in only 2.83 pixels of error in evaluation, indicating a successful training (Supplementary Figure 6A). Next, we ran the trained DLC model on the 4 training videos to get the frame-wise positions of the 6 body parts (~15 minutes). These outputs were then fed into B-SOiD to train a B-SOiD model for behavior classification (~10 minutes). B-SOiD uses unsupervised approach to form behavior groups and outputs video examples for each group, which is a convenient way for users to associate these computer-decided groups with ethologically meaningful behaviors (~5 minutes). We tested different number of behavior groups from ~100 to 5 groups and inspected the video examples for each group. We found that the 5-group were the best fit since: (1) others resulted in either too many different groups that turned out to be the same behavior that was expressed in different ways, or a single group containing different behaviors that could not be assigned to a single ethologically meaningful behavior type; and (2) the 5-group B-SOiD model showed 5 well-separated clusters (Supplementary Figure 6B). We then labeled each group as a single behavior type based on the most frequently observed behavior among the videos in that group (Supplementary Figure 6C). Finally, we ran the trained DLC model on the 3 testing videos (3 minutes in total at 30 fps) to produce the frame-wise body part position output and fed that into the trained B-SOiD model for behavior classification and quantification (~5 minutes).

*LabGym* does not require labeling or training to track the animals; users only need to specify a time window in a video for background extraction. Thus, as described above, we input the 4 training videos into *LabGym* to generate data pairs containing animations of user-defined duration and their paired pattern images (~5 minutes). We then sorted these data pairs into different folders according to user-defined behaviors of interest (~30 minutes). Next, we input these folders into *LabGym* to train a Categorizer for behavior classification (~10 minutes). Finally, we ran *LabGym* using the trained Categorizer on the 3 testing videos for behavior classification and quantification (~5 minutes).

The total time to set up and run DLC+B-SOiD on 8-minute training videos and 3-minute testing videos at 30 fps was about 9 hours (including 0.5 hour of labor), while that for *LabGym* was less than 1 hour (including 0.5 hour of labor; Figure 9B). To establish the ‘ground truth’ for determining the end accuracy of frame-wise identification of behaviors in the 3 testing videos, we manually annotated the behavior types in each frame of the 3 testing videos (1,800 frames per video). We then compared the frame-wise behavior classification outputs of B-SOiD or *LabGym* with this ground truth. We found that both B-SOiD and *LabGym* accurately identified rearing and locomotion in the ‘Base’ video (‘Amphetamine (baseline)’; Figure 9C and Supplementary Video 15). However, B-SOiD failed to identify head movements in ‘Post’ video (‘Amphetamine (post treatment)’), whereas *LabGym* achieved good accuracy in identifying this behavior (Figure 9D and Supplementary Video 16).

Both B-SOiD and *LabGym* provided quantitative measurements of the animal body kinematics during locomotion (Figures 9E and 9F), but only *LabGym* captured the real quantitative differences in behavior between rats given a single vs. repeated amphetamine exposure (Figure 9F). Specifically, after repeated exposure the frequency of locomotion was dramatically decreased while repeated head movements (i.e., in-placed stereotyped head movements) were increased (Supplementary Videos 15 and 16), causing a reduction in locomotion distance. This is a classic pattern that typifies the expression of psychomotor sensitization (Carr et al., 2020; Ferrario *et al.*, 2005). Additionally, it is noteworthy that the speed of locomotion was similar across groups. This highlights the need to quantify a behavior to accurately measure effects of psychostimulant drugs on behavior (see (Ferrario *et al.*, 2005) for additional discussion).

When the testing was applied to a video made in a different enclosure, the Categorizer in *LabGym*, which was trained on only 49~119 training data pairs for each behavior type, was able to generalize the behavioral diction with only a modest drop in accuracy (Figure 9G and Supplementary Video 17). In contrast, B-SOiD was not able to generalize well (Figure 9G).

It is possible that B-SOiD may be able to accurately identify the “head movement” behavior in the testing videos if additional training data were provided. Alternatively, it is possible that the pre-defined features (body length, angle, and speed) used by B-SOiD to form behavioral groups might not be useful for distinguishing head movements from other behaviors. Nevertheless, the purpose here was to directly compare approaches using the same training resource. Both approaches were able to accurately track rearing, but only *LabGym* was able to generalize to another testing enclosure, and capture behaviors specific to psychostimulant drug effects.

## DISCUSSION

*LabGym* is a unique computational tool that enables a generalizable solution to automated behavioral analysis. The flexibility and trainability of this tool allows it to be applied to very broad variety of species, behaviors and questions, and to be iteratively refined towards a particular set of conditions over time. This tool opens opportunities for automatic, high-throughput, targeted and customizable analyses of behaviors without the cost of commercial products, which will greatly facilitate the process of mechanistic discoveries in biological research. The standardized, visualizable behavioral datasets generated by *LabGym* will also benefit research in neuroscience, behavioral science, and computer science by helping to standardize procedures within and across research groups, and by providing long-term references for future studies to build on.

### Innovations in *LabGym*

Firstly, we developed a new method for effective background subtraction in *LabGym* that outperforms the state-of-the-art background subtraction approaches. This new method enables accurate tracking of animal bodies without the need of manual labeling and model training, which significantly saves the human labor and accelerates the analysis.

Secondly, unlike other tools using fixed and complex neural networks (Bohnslav *et al.*, 2021; Mathis *et al.*, 2018) that are slow to train and hard to generalize, we designed a new neural network structure in *LabGym* with customizable combinations of Animation Analyzer, Pattern Recognizer and Decision Maker. This allows users to customize the behavior categorization to suit behavior data with different complexity and enables fast running speed in even common laptops without GPUs. This new design also ensures the effectiveness in holistic assessment on both the complete spatiotemporal details in a behavior and the overall movement pattern that is unique to the behavior. The holistic assessment on behavioral information in *LabGym* does not rely on pre-defined features and does not require intermediate steps of extracting features, which minimizes the information loss.

Thirdly, we developed a new method for evaluating the overall pattern of the animal’s movement (including both the whole body and limbs) during a behavior. We designed the “pattern image” to exhibit the animal’s motion pattern of both the whole body and body parts efficiently and precisely. We then designed the Pattern Recognizer in Categorizer to analyze the pattern image. This method is more efficient and precise than current approaches for assessing motion flows that operate on every pixel in video frames to generate optical flows (Bohnslav *et al.*, 2021).

Lastly, *LabGym* is “end-to-end”: it provides a GUI and does not need intermediate or post-hoc steps that require users to have expertise in computation or programming. Instead, users only need to recognize and sort the animations and their paired pattern images as examples of a given behavior to ‘teach’ *LabGym* to identify the behavior. The behavior is then automatically quantified by *LabGym*.

### *LabGym* provides a way for users to generate standardized, visualizable behavioral datasets

In behavioral categorization based on deep learning approaches, the accuracy depends on the quality of training datasets, which should be diverse and well labeled (LeCun *et al.*, 2015; Roh *et al.*, 2021; Schmidhuber, 2015). In the computer vision field, high-quality datasets, such as ImageNet (Deng et al., 2009), have been established to facilitate the training of deep neural networks and to benchmark different algorithms. In neuroscience, however, such benchmark datasets for behaviors in model organisms are largely absent. A source of this problem is the lack of tools that can help researchers to generate standardized, visualizable behavioral datasets that are easy to label and to share. Although transfer learning that applies acquired network weights to new categorization problems can potentially be used to compensate for the lack of species-specific behavioral datasets, its efficiency relies on the similarity between existing datasets used for pretraining the networks and the new data (Yosinski et al., 2014). Current tools using transfer learning for behavioral categorizations (Bohnslav *et al.*, 2021) utilized network weights that were acquired from human behavioral datasets such as Kinetics-700 (Carreira et al., 2019). However, human behaviors are intrinsically different from the behaviors of model organisms, especially the soft-bodied ones. Therefore, species-specific behavioral datasets are still in critical demand for accurate behavioral categorizations in model organisms.

*LabGym* addresses the lack of such benchmark datasets by providing a way for users to generate standardized, visualizable behavioral datasets that to compare their categorization (labeling) of behavior and refine the labeling. Importantly, such expert-labeled behavioral datasets, if shared in the research community, not only ensure the accurate and reproducible executions of categorization criteria across different experimenters or trials, but also provide benchmarks for comparing different algorithms, and references for future studies to build on.

### Limitations of *LabGym*

*LabGym* does not require labeling or training to track the animals. In turn, it does have preferred settings for video recording. Although *LabGym* does not have restriction on the kind of background/arena/camera to use, the background/arena/camera needs to be static during the period for behavior analysis. The illumination can have dark-to-bright or bright to dark transitions but need to be stable before and after the transitions during the period for behavior analysis. Animals are expected to present some locational changes during video recording, instead of being immobile all the time. The time window during which animals are moving can be used for background extraction in *LabGym*. This requirement in video recording can be alleviated by implementing mask region-based convolutional neural networks (mask R-CNN) (He et al., 2017) in future versions of *LabGym*. Mask R-CNN can be used to segment animals from dynamic backgrounds (i.e., the backgrounds continually change over time). Currently, when analyzing animals in dynamic backgrounds, users may first consider using the mask R-CNN (https://github.com/matterport/Mask_RCNN) (Abdulla, 2017) to obtain animal-masked videos, and then use the current version of *LabGym* to analyze such videos. Additional labeling and training efforts will be needed in such cases.

The current version of *LabGym* is not optimized for analyzing social behaviors. In social behaviors, animals have body contacts. The animal identity reassignment after animals re-separate might not be accurate in *LabGym*. The accuracy of animal identity reassignment can be improved by implementing mask R-CNN in future versions of *LabGym*.

## ACKNOWLEDGMENTS

We thank Drs. Daniel Leventhal, Hanchuan Peng, and Guoqiang Yu for helpful comments on earlier versions of the manuscript. This research was supported by National Institutes of Health (NIH) to B.Y. (R01NS104299 and R01EB028159), and C.R.F. (R01-DK106188, R01-DK115526, R01-DA044204). *Drosophila* Stocks from the Bloomington *Drosophila* Stock Center (NIH P40OD018537) were used in this study. The content is solely the responsibility of the authors and does not necessarily represent the official views of the NIH.

## AUTHOR CONTRIBUTIONS

Y.H. and B.Y. conceived the project. Y.H. designed and programmed *LabGym* with the guidance of J.Z. and B.Y.. Y.H., K.I., H.W., C.R.F., A.G., and B.W. performed the behavioral experiments in *Drosophila* larvae (Y.H, K.I. and H.W.), rats (C.R.F.), and mice (A.G. and B.W.). Y.H., K.I., H.W., and Y.X. generated and labelled the dataset for larval behaviors (Y.H., K.I., H.W. and Y.X.), rat behaviors (Y.H.), and mouse behaviors (Y.H. and A.G.). Y.H. performed the computational analyses and the validation of behavioral categorizations with the assistance from C.R.F., A.G., B.W., K.I., and H.W.. Y.H, A.B.M. and R.B.I. performed the benchmark comparison between *LabGym* and DeepLabCut+B-SOiD. Y.H., C.R.F., B.W., J.Z., and B.Y. wrote the manuscript.

