## Supplementary information for "*LabGym*: quantification of user-defined animal behaviors using learning-based holistic assessment"

**Inventory of Supplemental Information**

Resource availability

Methods

References for supplemental information

Supplementary Figures 1-6.

Supplementary Tables 1 and 2.

Supplementary Videos S1 to S17.

**METHORDS**

**Data and Code Availability**

The behavioral datasets generated and analyzed in the current study for *Drosophila* larvae, mice and rats will be deposited for free access. The open-source code of *LabGym* is available in GitHub (http://github.com/umyelab/LabGym).

**Overall implementation**

*LabGym* was written in Python programming language (version 3.9) (Van Rossum and Drake, 2000). Python libraries used in *LabGym* are: Numpy (Harris et al., 2020), Scipy (Virtanen et al., 2020), scikit-learn (Pedregosa et al., 2011), scikit-image (van der Walt et al., 2014), MoviePy, Matplotlib (Hunter, 2007), OpenCV (Bradski, 2000), Pandas (McKinney, 2010), Seaborn (Waskom et al., 2017), Tensorflow (Abadi et al., 2016), wxPython (Rappin and Dunn, 2006).

**Computation hardware**

The computational procedures in this study were performed on the following 4 computers. (1) a MacBook Pro 13 inch (early 2015 model) with Mac OS Big Sur (or later), 3.1 GHz Dual-Core Intel Core i7 processor, 16 GB 1867 MHz DDR3 memory, and Intel Iris Graphics 6100 1536 MB graphics. (2) HP Z4 Workstation with Windows 10 Enterprise for Workstations, 3.2 GHz Intel Xeon W-2104 processor, 32GB 2666 MHz DDR4, and NVIDIA GeForce GTX 1080 Ti. (3) HP Z4 G4 Workstation with Windows 10 Pro for Workstations, 3.6 GHz Intel Xeon W-2133 processor, 32GB 2666 MHz DDR4, and NVIDIA Quadro P2000. (4) DESKTOP-G2HRLHT with Windows 10 Pro for Workstations, 3.6 GHz Intel Core i9-9900K CPU, 64 GB 2666 MHz DDR4, and NVIDIA GeForce RTX 2080.

**Remove backgrounds and track multiple animals**

*Reconstruct the static backgrounds for a video*

To reconstruct the static background of a video, the stable intensity value over time for each pixel, denoted as $P_{stable,}$ was identified by looking for a sliding time window $t_{stable}$ of 100 frames (about 3 sec if the fps is 30) during which the value of this pixel was stable. $P_{stable}$ is the mean intensity during $t_{stable}$, which is obtained by searching through the entire duration of the video. Specifically, for a video in which the animals were generally lighter than the background, it was identified when the sum of the mean and standard deviation of pixel values in the time window was the smallest (Equation 1). For a video in which the animals were generally darker than the background, $t_{stable}$ was the time window in which the sum of the mean and standard deviation of the inverted pixel values was the smallest (Equation 1).

$$P_{stable} = \overline{P(t_{stable})},$$
$$t_{stable} = \begin{cases} \underset{t}{argmin}(\overline{P(t)} + std(t)) & if\ body\ part\ lighter\ than\ background \\ \underset{t}{argmin}((255 - \overline{P(t)}) + std(t)) & if\ body\ part\ darker\ than\ background \end{cases} \quad (1)$$

where *t* represents 100-frame time windows sliding over the video, *std(t)* is the standard deviation of the pixel values during the time window *t*.

In the less likely case that the animals were partially lighter or darker than the background, $P_{stable}$ was the mean of this pixel over the entire duration of the video. In addition, if the illumination was generally stable in the entire video, then we simply used the extremum value (the minimum for animals lighter than the background, and the maximum for animals darker than the background) to achieve similar results as using $P_{stable}$ while keeping the computational cost low. Therefore, we provide users with the option to specify whether the illumination in the video is stable.

The $P_{stable}$ for the time when the illumination changes (for example, the optogenetic manipulation period) was identified separately when the mean value of all the pixels in the current frame was 1.2-fold greater than that in the first frame.

*Track the animals*

The foreground (all the pixels of an animal) was obtained by subtracting background in each frame. The approximate area of single animal was estimated by averaging the total area of all the animals over the animal number. Each animal is a matrix of [identity, [contours], [centers], [behavioral information], …]. We assumed that the Euclidean distance between the centers of the same animal in consecutive frames is always the smallest among the distances between all the center pairs. In this way, all the animals in current frames are linked to the same animals in previous frames and the new information of the same animals is added into the animal matrices. Since the animals might be undetectable in some frames, we examined the entire frames from the beginning to the end of the analysis, rather than only examine the consecutive frames, to minimize the probability of not being tracked.

Multiple animals in the same enclosure sometimes collide with each other, which might cause false tracking. To make the analysis fully automatic, *LabGym* excludes animals from tracking if they do not meet the user-defined criteria for the number of entangled animals to be allowed for analysis. For example, if entanglement is not allowed, the merged foreground of two or more entangled animal would be temporarily excluded from tracking until they separate again. The animal that is lost track for over one second will be reassigned by a new ID/matrix and the previous ID/matrix for this animal will be deactivated. Any ID/matrix that is deactivated for longer than 20% of the entire analysis duration is considered as disappeared and is automatically excluded from the analysis, which ensures that the analysis is only performed on the reliably tracked animals. When the entanglement is allowed, merged foreground will always be tracked and animal IDs/matrices can be preserved when they separate again.

**Generation of animations and pattern images**

*Generate animations*

The animation of an animal is a set of blobs over a user-defined time window. Each blob is a cropped video frame containing a single animal. The animal foreground in a blob is masked by the animal contour and the area inside the contour in the original frame and the background is set to pixel value of 0. Users also have the option to include backgrounds in the animations since the environmental factors sometimes are key to behavioral categorization.

*Generate pattern images*

The pattern image for each animation was generated by drawing all animal contours (the whole body and the body parts) within the animation onto a black background with gradual changing colors to indicate the temporal sequences of the contours. The sizes of pattern images are the same as the frame in their paired animations. Users can also choose whether to show animals' body parts in the pattern images, depending on their needs.

**The design and implementation of Categorizer**

*Animation Analyzer*

Animation Analyzer consists of convolutional blocks warped with time distributed layers and followed by recurrent layers (long short-term memory, LSTM) (Abadi *et al.*, 2016; Fukushima, 1980; Hochreiter and Schmidhuber, 1997). It takes 4-dimentional (4-D) tensors (timestep, width, height, color channel) as inputs, in which the timestep, width/height and color channel are all user definable. The timestep of the input tensors is the number of frames in an animation (the length of an animation). Ideally the length should precisely match the variable durations of different behaviors so that each animation only contains a single behavior. However, since deep neural networks require their input shape to be consistent during the analysis, the durations for all the animations need to be the same. We did not use zero paddings to arbitrarily make the input shape consistent because we wanted the Animation Analyzer to learn to ignore those frames mixed with behaviors of nontargeted categories, which is a practical scenario in behavioral analyses. To achieve the best efficiency in training the Categorizer, the duration of the animations should be the shortest time for experimenters to distinguish all the behavioral categories. Animation Analyzer implements two different architectures: VGG-like (Simonyan and Zisserman, 2014) and ResNet-like (He et al., 2016). There are 7 different complexity levels of Animation Analyzer for user to choose to fit different datasets. Levels 1, 2, 3 or 4 is VGG-like architecture with 2, 5, 9, or 13 convolutional layers (Conv2D), respectively. Each convolutional layer is followed by a batch normalization layer (BN). Max pooling layers (MaxPooling2D) are added after the $2^{nd}$ BN in level 1, after the $2^{nd}$ and $5^{th}$ BN in level 2, after the $2^{nd}$, $5^{th}$ and $9^{th}$ BN in level 3, or after the $2^{nd}$, $5^{th}$, $9^{th}$ and $13^{th}$ BN in level 4. Levels 5, 6, or 7 is ResNet18, ResNet34 or ResNet50 architecture. After the convolutional operations, the outputs are flattened into 1-dimentional (1-D) vectors at each time step and then are passed to LSTM. The outputs of LSTM are passed to the Decision Maker submodule.

To make *LabGym* applicable to behavioral datasets of various complexities, the frame sizes and color channels of the input tensors are user definable. Therefore, the network architectures in Animation Analyzer are provided with various options, from simple 2-layer VGG-like to complex 50-layer ResNet50 architectures. The number of filters in the first convolution layer is determined by the height or width dimension of input tensors (the height and width of the input tensor is the same), and then doubled after each max pooling layer. The number of filters in the LSTM is the same as that of the last convolution layer.

*Pattern Recognizer*

Pattern Recognizer consists of convolutional blocks and takes 3-dimentional (3-D) tensors (width, height, color channel) as inputs, in which both the width and height are user definable. The color channel is fixed to 3 (Red, Green, and Blue; RGB) since the colors in the pattern images indicate the temporal sequences of animal motions. There are also 7 different complex levels of Pattern Recognizer for the user to choose from to suit different datasets. The architecture for each level in Pattern Recognizer is the same as that for each level in Animation Analyzer, except that there is no time-distributed wrapper in Pattern Recognizer.

*Decision Maker*

Decision Maker consists of a concatenation layer and two fully connected (dense) layers. The concatenation layer merges the outputs from both Animation Analyzer and Pattern Recognizer into a single 1-D vector and outputs to the first dense layer. The first dense layer then outputs to the second dense layer in which a SoftMax function is used for computing the probabilities of all the behavioral categories. The number of nodes in the first dense layer of Decision Maker is determined by the complexity levels of Animation Analyzer or Pattern Recognizer, whichever is higher. This association allows the Decision Maker to adapt based on the structure of the other two modules. A Batch Normalization layer and a Dropout layer (dropout rate is set to 0.5) are added before the second dense layer. The number of nodes in the second dense layer is set to the number of behavioral categories. At each frame during the analysis, the output of Decision Maker for an animal is a matrix of probabilities for all the behavioral categories and the behavioral category with the highest probability is determined to be the behavior that the animal performs. Users have the option of setting a threshold to make *LabGym* output an 'Uncertain' decision if the difference between the highest and the second highest probabilities is smaller than the threshold. If the behavioral category is determined as 'Uncertain', an 'N/A' will be shown for the animal at the frame in the annotated video and the block for the animal at the frame in the raster plot will be white (no color). The probabilities for all the behaviors at each frame are exported to Microsoft Excel files for the usage as users see fit.

**Train Categorizers**

The loss function used to train Categorizers is either binary crossentropy if the number of behavioral categories is two, or categorical crossentropy if the number of behavioral categories is three or more. Stochastic gradient descent (SGD) (Ruder, 2016) with an initial learning rate of $1 \times 10^{-4}$ is used for optimizer in training. The learning rate decreases by a factor of 0.2 if the validation loss stops decreasing (decreasing by <0.001 is considered as 'stop decreasing') for 2 training epochs, and the training stops if the validation loss stops decreasing for 4 training epochs. The model is automatically updated after a training epoch reaches a minimal validation loss. The batch size in training is 8, 16, or 32 if the number of validation data is less than 5,000, between 5,000 and 50,000, or over 50,000, respectively. The ratio of training and validation data split is 0.8:0.2.

To enhance the training efficacy and the generalizability of the trained models, we developed 9 different data augmentation methods that apply the same random alterations to both an animation and its paired pattern image. Notably, during data augmentation, all the alterations of frames/images are performed only on animal foregrounds and their contours. The absolute dimensions of frames/images and the black backgrounds are unchanged. These methods are performed in combination and users have the options to choose which method to perform. If all the data augmentation methods are applied, the amount of data can be expanded to 47 times of its original amount.

1. Random rotation: both the animations and their paired pattern images are rotated with a random angle in ranges between 10°~50°, 50°~90°, 90°~130°, 130°~170°, 30°~80° and 100°~150° (the latter two ranges are used to be combined with random brightness changes).
2. Horizontal flipping: both the animations and their paired pattern images are flipped horizontally.
3. Vertical flipping: both the animations and their paired pattern images are flipped vertically.
4. Random brightening: the animations have brightness increase in ranges between 30~80.
5. Random dimming: the animations have brightness decrease in ranges between 30~80.
6. Random shearing: both the animations and their paired pattern images are sheared with a random factor in ranges between -0.21~-0.15 and 0.15~0.21.
7. Radom rescaling: both the animations and their paired pattern images are rescaled in either their widths or heights with a random ratio in a range between 0.6~0.9.
8. Random deletion: one or two frames in the animations will be randomly deleted and replaced with black images of the same dimensions.
9. Exclude original: exclude the original data, which can be used to retrain a network on the same dataset.

**Select suitable Categorizers for different behavioral datasets (Supplementary Table. 1)**

The Categorizer can be customized into different complexity to address behavioral datasets with different complexity. The complexity of Categorizer is determined by two factors: the level of Animation Analyzer / Pattern Recognizer and the size of input frame / image. The former determines how deep whereas the latter determines how wide the Categorizer is (see **The design and implementation of Categorizer**). There are 7 levels of Animation Analyzer / Pattern Recognizer for user to choose and the input frame size is completely defined by users. The general principle to guide the selection of suitable Categorizer for each behavioral dataset is Occam's razor, which means that if the performances of two Categorizers are comparable, the simpler one is better. Therefore, we started from Categorizers with the simple complexity and gradually increased the complexity until their performance are satisfying, for all 3 behavioral datasets.

The behaviors in the larva dataset are distinguishable from the sequential changes of their body shapes during the behaviors. Besides, the resolution in the animations is insufficient to show details of the larva body. Therefore, we chose to downsize the frame in animations into 8 x 8 (gray scale) for Animation Analyzer so that it can focus on only the body shapes without learning too many irrelevant details. The pattern images for larva behaviors contain more details such as the lines of different colors indicating the temporal sequences of the behaviors. Therefore, we chose 32 x 32 (RGB scale) as the input image size of Pattern Recognizer. We started from the level 1 of Animation Analyzer and level 2 of Pattern Recognizer, which already achieved excellent overall accuracy on the larva nociceptive behavior subset. When we increased the complexity of the Categorizer to level 1 Animation Analyzer and level 3 Pattern Recognizer, the accuracy decreased, indicating the potential overfitting of the Categorizer. Therefore, we stopped at level 1 Animation Analyzer and level 2 Pattern Recognizer as the suitable Categorizer for the larva dataset (*LarvaN*).

To select the suitable Categorizer for the rat dataset, we chose 16 x 16 or 32 x 32 (gray scale) for the input frame size of Animation Analyzer and 32 x 32 or 64 x 64 (RGB scale) for the input image size of Pattern Recognizer. This was because the differences among different behaviors in the rat dataset are more subtle than those in the larva dataset and we wanted the Categorizer to learn more details. The overall validation accuracy stopped increasing at level 4 / 4 for Animation Analyzer / Pattern Recognizer but was still not ideal (0.8), indicating that the unsatisfying accuracy was not because of low complexity of the Categorizer. Note that due to the data-driven nature of deep-learning based tools, the accuracy of Categorizer also largely relies on the quality of the training dataset such as the amount and diversity of the data, and the labeling accuracy. Therefore, we refined the rat dataset by discarding some examples that were ambiguous for categorization by the labeler. We then trained the Categorizer of level 4 / 4 for Animation Analyzer (32 x 32 x 1) / Pattern Recognizer (64 x 64 x3) on the refined rat dataset, which achieved significantly improved overall accuracy and was determined as the suitable Categorizer for the rat dataset (*RatA*).

To select the suitable Categorizer for the mouse dataset, we included all the categories in which the number of examples (the number of animations and pattern images in pair) is greater than 50. Since the animations in the mouse dataset contain details that are key to categorizations of the behaviors, such as the movement of paws / noses, we chose 32 x 32 or 64 x 64 (gray scale) for Animation Analyzer and 32 x 32 or 64 x 64 (RGB scale) for Pattern Recognizer. We found that level 4 / 4 for Analyzer (64 x 64 x 1) / Pattern Recognizer (64 x 64 x3) achieved satisfying overall accuracy and determined it as the suitable Categorizer for the mouse dataset (*MouseH*).

All selected Categorizers were re-trained once on the data augmented from the same original datasets (with the original data excluded).

**Test Categorizer in practice**

*LabGym* provides two convenient ways of validation on how Categorizers perform. First, the copies of behavioral videos with full annotations of the behaviors assist the real-time visualization of both the tracking and the behavioral categorizations. Second, for a more solid test, users can use Generate Datasets functional unit in the GUI of *LabGym* to generate unsorted behavioral data (an animation and its paired pattern image) and sort them into different categories (into folders under the behavior names) to build a ground-truth testing dataset. Users can then use this ground-truth testing dataset in Test Networks functional unit to do validations. In this way, the users are blinded to the predictions of the Categorizer to be tested and thus the validation is unbiased. In this study, all tests for the in-practice performance of the *LarvaN*, *RatA* and *MouseH* were performed through this approach.

We used the following performance metrics in testing Categorizers: precision (equation 2), recall (equation 3), f1 score (equation 4), overall accuracy (equation 5).

$$precision_i = \frac{true\ positives_i}{true\ positives_i + false\ positives_i} \quad (2)$$

$$recall_i = \frac{true\ positives_i}{true\ positives_i + false\ negatives_i} \quad (3)$$

$$f1\ score_i = \frac{2 \times (precision_i \times recall_i)}{precision_i + recall_i} \quad (4)$$

$$overall\ accuracy = \frac{total\ correct\ predictions}{total\ predictions} \quad (5)$$

**Calculate behavioral parameters**

In the current version of *LabGym*, 14 behavioral parameters are computed based on the information of behavioral categories and animal foregrounds: acceleration / velocity reduction, angle, count, distance, duration, intensity (area), intensity (length), latency, magnitude (area), magnitude (length), speed, velocity, vigor (area) and vigor (length).

1. The angle is the movement direction (against to the animal body axis) of the animal during a behavior, which is the mean of all the included angle ($\theta$) between animal body axis and the movement direction during the time window (*t*) for categorizing the behavior (equation 6).

$$angle = \frac{\sum_{i=n-t}^{n}(\theta_i)}{\text{n}} \qquad (6)$$

2. The count is the summary of the behavioral frequencies, which is the occurrence number of a behavior within the entire duration of analysis. Consecutive single occurrences (at a single frame) of the same behavior are considered as one count.
3. The distance is the total distance traveled of the animal by performing a behavior within the entire duration of analysis.
4. The duration is the summary of how persistent a behavior is, which is the total time of a behavior within the entire duration of analysis.
5. The intensity (area) / intensity (length) is the summary of how intense a behavior is, which is the accumulated proportional changes of the animal body area (*a*) / length (*l*) between frames divided by the time window for categorizing the behaviors (*t*) when performing a behavior (equations 7 and 8).

$$intensity\ (a) = \frac{a}{t}\ ,$$

$$a = \sum_{i=0}^{n}\left(\frac{a_n - a_i}{a_i}\right) \qquad (7)$$

$$intensity\ (l) = \frac{l}{t}\ ,$$

$$l = \sum_{i=0}^{n}\left(\frac{l_n - l_i}{l_i}\right) \qquad (8)$$

6. The latency is the summary of how soon a behavior starts, which is the time starting from the beginning of the analysis to the time point that the behavior occurs for the first time.
7. The magnitude (area) / magnitude (length) is the summary of the motion magnitude, which is the maximum proportional change in animal body area (*a*) or length (*l*) when performing a behavior (equations 9 and 10).

$$magnitude\ (a) = \max_{0 \le i \le n}\left(\frac{a_n - a_i}{a_i}\right) \qquad (9)$$

$$magnitude\ (l) = \max_{0 \le i \le n}\left(\frac{l_n - l_i}{l_i}\right) \qquad (10)$$

8. The speed is the summary of how fast the animal moves when performing a behavior, which is the total distance traveled (*d*) (between the two centers of mass of the animal) during the time window for categorizing the behavior divided by the time window (equation 11).

$$speed = \frac{\sum_{i=n-t}^{n}(d_i)}{t} \qquad (11)$$

9. The velocity is the summary of how efficient the animal's movement is when performing a behavior, which is the maximum displacement (*dt*) (between the two centers of mass of the animal) divided by the time ($t_{occuring}$) that such displacement takes place (equation 12).

$$velocity = \frac{\max_{0 \le i \le n} dt_i}{t_{occuring}} \qquad (12)$$

10. The acceleration / velocity reduction is the summary of how fast the animal's velocity changes while performing a behavior, which is the difference between maximum velocity ($v_{max}$) and minimum velocity ($v_{min}$) divided by the time ($t_{occuring}$) that such velocity change takes place (equation 13).

$$acceleration = \frac{v_{max} - v_{min}}{t_{occuring}} \qquad (13)$$

11. The vigor (area) / vigor (length) is the summary of how vigorous a behavior is, which is the magnitude (area) / magnitude (length) divided by the time ($t_{occuring}$) that such a change takes place (equations 14 and 15).

$$vigor\ (a) = \frac{magnitude\ (a)}{t_{occuring}} \qquad (14)$$

$$vigor\ (l) = \frac{magnitude\ (l)}{t_{occuring}} \qquad (15)$$

**Criteria for labeling the behavioral categories**

*Drosophila larva*

*Crawling*: A larva performs one of the following actions: 1) peristalsis 2) moving with body mostly straight 3) moving along the direction mostly aligned with the body axis. The crawling results in positional changes of larval body that can be clearly seen in the pattern image.

*Curling*: A larva performs ‘C’-shape bending and both the head and tail bend approximately at the same time from a straight, resting body shape.

*Hunching*: A larva retracts only the head.

*Rolling*: A larva moves with a curling body shape along the anterior-posterior body axis, which causes lateral positional changes in the larval body that can be seen in the pattern image.

*Turning*: A larva performs a body bending that is not in ‘C’-shape (body bending that is not curling).

*Uncoiling*: A larva performs the actions in temporal sequences that are opposite to curling or turning (from a coiling body shape to a straight one).

*Immobile*: A larva does not move or only has subtle movements.

*Mouse*

*Behind the wheel*: A mouse is positioned behind the running wheel.

*Body grooming*: A mouse licks its body, below the neck.

*Chewing*: A mouse chews on food pellets.

*Coming down*: A mouse lowers from a standing pose on its hindpaws to a sitting pose on both forepaws and hindpaws.

*Crawling*: A mouse crawls under the running wheel.

*Face grooming*: A mouse uses its paws to groom the face, head, mouth and/or ear.

*Foraging*: A mouse walks with its snout on the cage bedding, digs through the cage bedding with its forepaws, or investigates the cage bedding with its forepaws while stationary.

*Hind paw grooming*: A mouse uses its hindpaws to groom its face or body.

*Jumping onto the wheel*: A mouse jumps onto the running wheel.

*Nest building*: A mouse uses its mouth to build a nest or reposition nestlet material in the homecage.

*Rearing up*: A mouse rises from a sitting position with both forepaws and hindpaws on the cage bedding, to a standing position with only the hindpaws on the cage bedding.

*Resting on the wheel*: A mouse remains stationary on the running wheel.

*Running on the wheel*: A mouse runs on the running wheel.

*Sleeping*: A mouse sleeps in the homecage.

*Sniffing*: A mouse repeatedly sniffs the air, with the snout repeating superior and inferior movements while pointed at the cage walls or the lid of the homecage.

*Standing*: A mouse remains in a standing posture on both hindpaws.

*Turning*: A mouse turns to show a frontal view, a side profile view, or a back view.

*Unknown bv*: A mouse is performing some behavior while only its back is visible to the camera. The behavior is unidentifiable, unless the mouse is rearing up, coming down, or standing.

*Walking*: A mouse walks on the bedding (not on the running wheel).

*Rat*

*Body grooming*: A rat grooms its body.

*Face grooming*: A rat grooms its face.

*Head swaying (moving)*: A rat sways its head side to side for more than one repeats.

*Locomotion*: A rat proceeds forward.

*Orientating*: A rat turns or repositions.

*Rearing*: A rat rears up.

*In place*: A rat rests in place with some small movements.

*Still*: A rat is completely motionless.

**Behavioral experiments**

*Drosophila larval behaviors responding to nociceptor stimulation*

Nociceptors of late 3$^{rd}$ instar larvae were activated optogenetically *TrpA1*-QF > QUAS-ChR2$^{T159C}$ (Bloomington *Drosophila* stock center #36345 and #52260) transgenes. To stimulate *TrpA1*-QF > QUAS-ChR2$^{T159C}$, 50 μW/mm$^2$ blue light was applied for 2.5 sec. GtACR1 was expressed in LK neurons to inhibit these neurons (Hu et al., 2020). GtACR1-mediated optogenetic inhibition was done with 2.5 sec of amber light (590 nm). All LEDs used were from LUXEON StarLED. Embryos were collected on standard cornmeal containing 500 μM all-trans-retinal (ATR) (A.G. Scientific).

For each trial in larval behaviors, we placed 10-15 larvae on a ∅ 35-mm or 90-mm plate covered with a thin layer of water. The larvae behaviors were recorded by using a webcam (Logitech) at a resolution of 1280 x 720 or 864 x 480 resolution (30 fps) in the MP4 or AVI format.

*Drosophila larval behaviors responding to sound stimuli*

Late 3$^{rd}$ instar larvae were exposed to white noise at 80 or 94 dB SPL (0.2 – 20 kHz) for 5 sec. The larvae were tested on a black agar plate to increase the contrast between the background and the foreground for a better signal-to-noise ratio. All larvae were raised in a custom-made sound-resistant box for 5-7 days at room temperature to minimize sound exposure, and were recorded with a Logitech 4K Brio webcam (30 fps) at 1080 x 1080 resolution in the MP4 format.

*Drosophila adult courtship behaviors*

To demonstrate the potential capacity of *LabGym* in analyzing social behaviors, we used some videos of *Drosophila* courtship behaviors from a previous report (Hu et al., 2014). In these experiments, a wild-type male (*Oregon R*) was paired with either one or 4 females (*Oregon R* or $w^{1118}$) in a mating chamber.

*Psychomotor Sensitization in Rats*

All rodent procedures were approved by University of Michigan Institutional Animal Care and Use Committee and are consistent with NIH guidelines.

Adult male Sprague Dawley rats (~55 days old at the start of the experiment) were pair-housed on reverse 12/12 hr light/dark cycle throughout, food and water were available ad libitum and rats were tested in red light conditions during the dark phase of the cycle (Envigo; Indianapolis, IN). All testing occurred in custom made chambers (13"W x 26.5"L x 24"T) equipped with overhead video cameras (fps:30; frame size: 928 x 480; video format: .MP4). Behavior was recorded throughout each session. Rats were first habituated to the testing chambers and injection procedure; they were placed in the chamber for 40 min, given an i.p. injection of saline (0.9%, 1 ml/kg) and returned to their home cage after an additional 40 min (4 sessions, 1/day). Rats were then split into two groups given i.p. injections of either saline (N=3) or d-amphetamine sulfate dissolved in saline (Sigma Aldrich, N=7). On each injection day rats were placed in the chambers for 40 min prior to injection. On the first day, responses to increasing doses of d-amphetamine were determined within session (0.32, 1.0, 3.2, and 5.6 mg/kg, i.p), as previously described (Robinson et al., 2015). The following day rats began a sensitization regimen in which they were given increasing concentrations of d-amphetamine (0.5, 2, 4, 5, 6, 0.5, 6, 6, 0.5 mg/kg, i.p.) with one injection given per day, adapted from (Ferrario and Robinson, 2007). The within session dose response was then re-assessed after 20-21 days of withdrawal. Rats in the saline group received saline throughout. The rat videos used for demonstration of tracking in *LabGym* were sample video in previous reports (Ellinwood and Balster, 1974; Ferrario et al., 2005).

*Mouse*

All rodent procedures were approved by University of Michigan Institutional Animal Care and Use Committee and are consistent with NIH guidelines.

Adult male C57BL/6J mice (one animal aged ~ 80 weeks old at the start of the video recordings, one animal aged ~32 weeks old at the start of the video recordings) were single housed in cages within a humidity and temperature-controlled vivarium and kept on a 12/12 hr light/dark cycle (lights on 6 am) with ad libitum access to food and water. At the time of video recording, both mice were single-housed in a cage with bedding, ¼ Nestlet (Ancare, Bellmore, NY) for nest building, and a Mouse Igloo and Fast Trac running wheel (Bio-Serv, Flemington, NJ), located within a custom-made white PVC foam sheet (American Plastics Solutions, Saline, MI) enclosure, located within a humidity and temperature-controlled room. Animals had ad libitum access to food and water during the video recordings and were kept on a 12/12 hour light/dark cycle (lights on 6 am). Behavior was recorded with a Basler acA1300-200um camera, equipped with a Basler C123-0418-5M lens and a NIR Bandpass Infrared Filter (Machine Vision Store, St. Paul, MN). InfraRed lighting (850 nm, LEDLightsWorld) diffused by a ¼" x 13.5" x 24" sheet of white acrylic plexiglass (estreetplastics, Royse City, Tx) was used to illuminate the homecage as well as to provide the background for all video recordings. Free home-cage behavior was recorded in four-hour segments as .mkv files, at a resolution of 1280 x 1024, 30 fps, with an exposure time of 6500 μs.

### Supplementary Figure 1

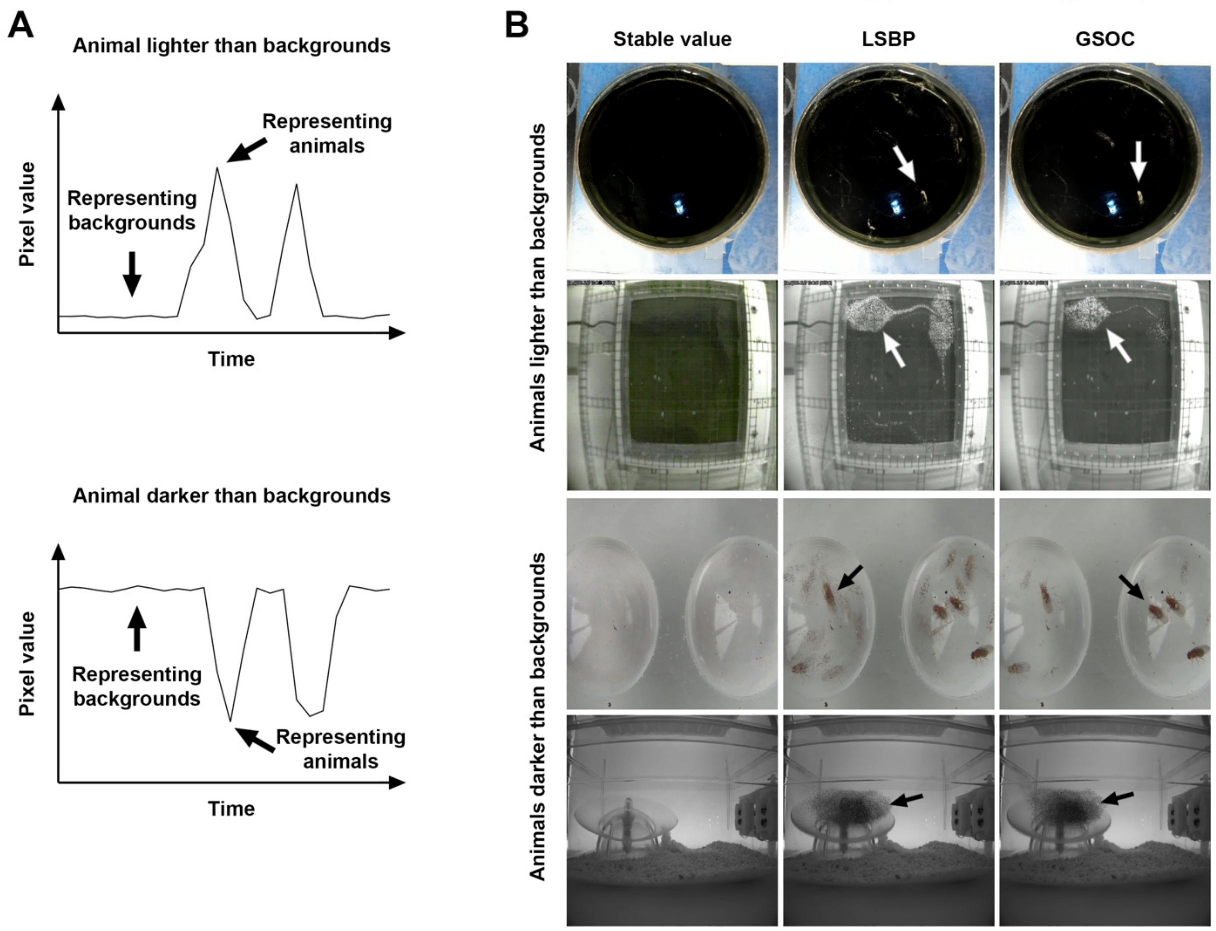


**Supplementary Figure 1. The stable-value detection method outperforms the state of the art.**

**A**. Illustrations showing the designing rationale of stable-value detection method for reconstructing the static background of a video in two different scenarios (animal lighter or darker than the backgrounds).

**B**. Examples for reconstructed static backgrounds of the same videos using different methods. Arrows point at the remaining animal traces in the reconstructed static backgrounds.

### Supplementary Figure 2

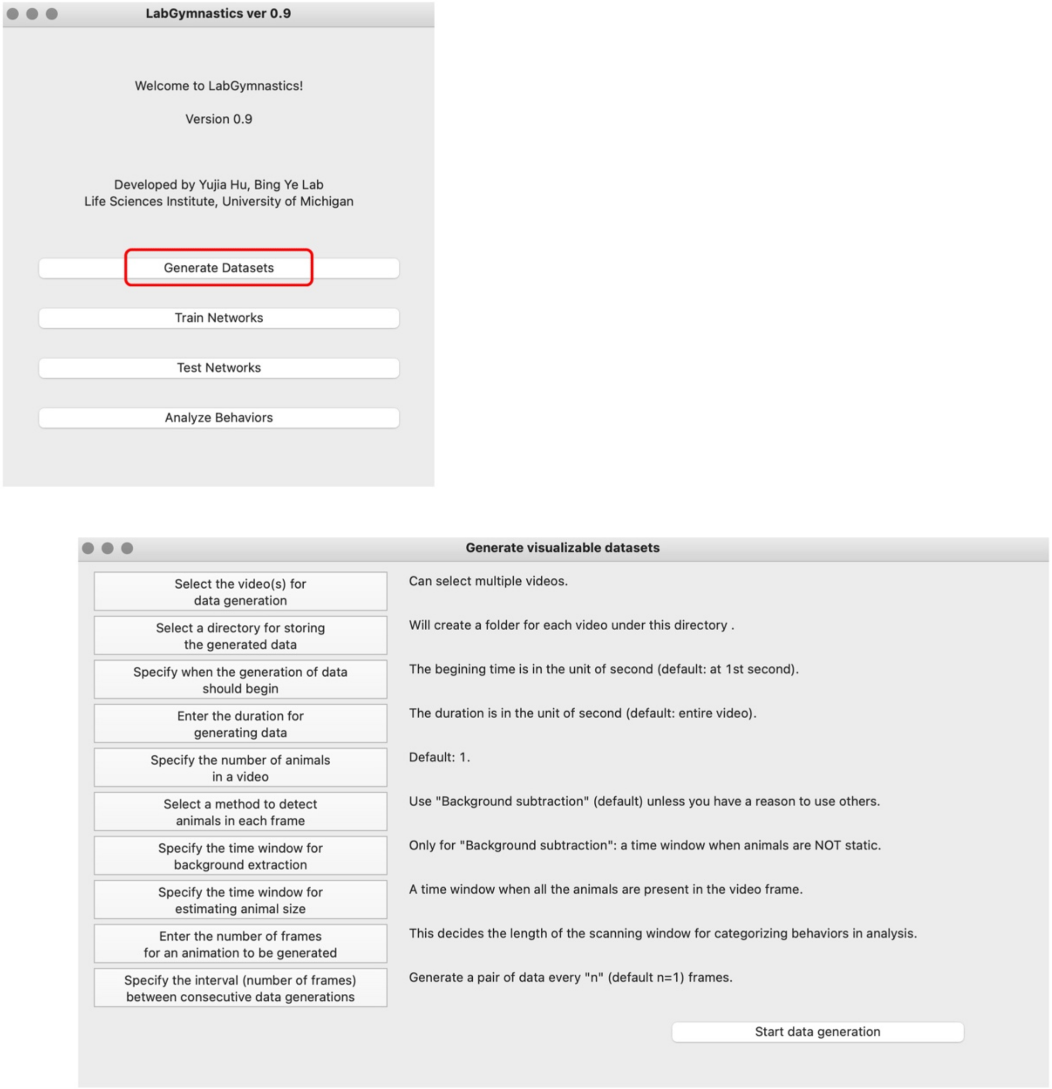


**Supplementary Figure 2. The graphical user interface (GUI) for the Generate Data functional unit.**

This unit offers options for batch processing of behavioral videos to generate standardized behavioral data (animations and their paired pattern images). The standardized data can be used to establish benchmark datasets for training the Categorizer. The functional buttons are on the left side; the short explanations and recommendations are on the right side.

### Supplementary Figure 3

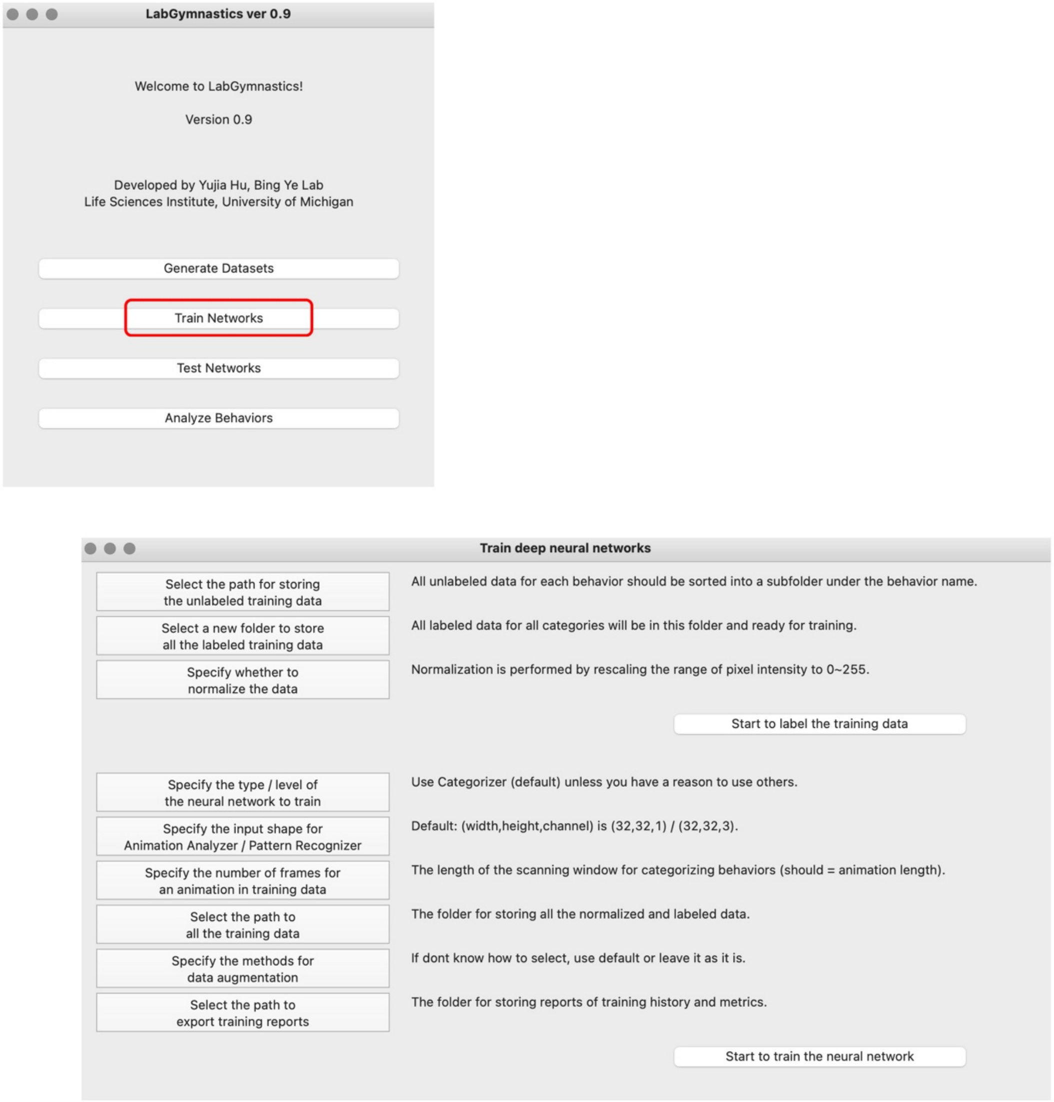


**Supplementary Figure 3. The GUI for Train Networks functional unit.**
This unit offers options for training Categorizer with various complexity. This unit helps to create user-defined labels (behavioral categories) and normalize the training data. After training, the trained Categorizers are added into the Analyze Behaviors unit for behavioral analysis. The functional buttons are on the left side; the short explanations and recommendations are on the right side.

### Supplementary Figure 4

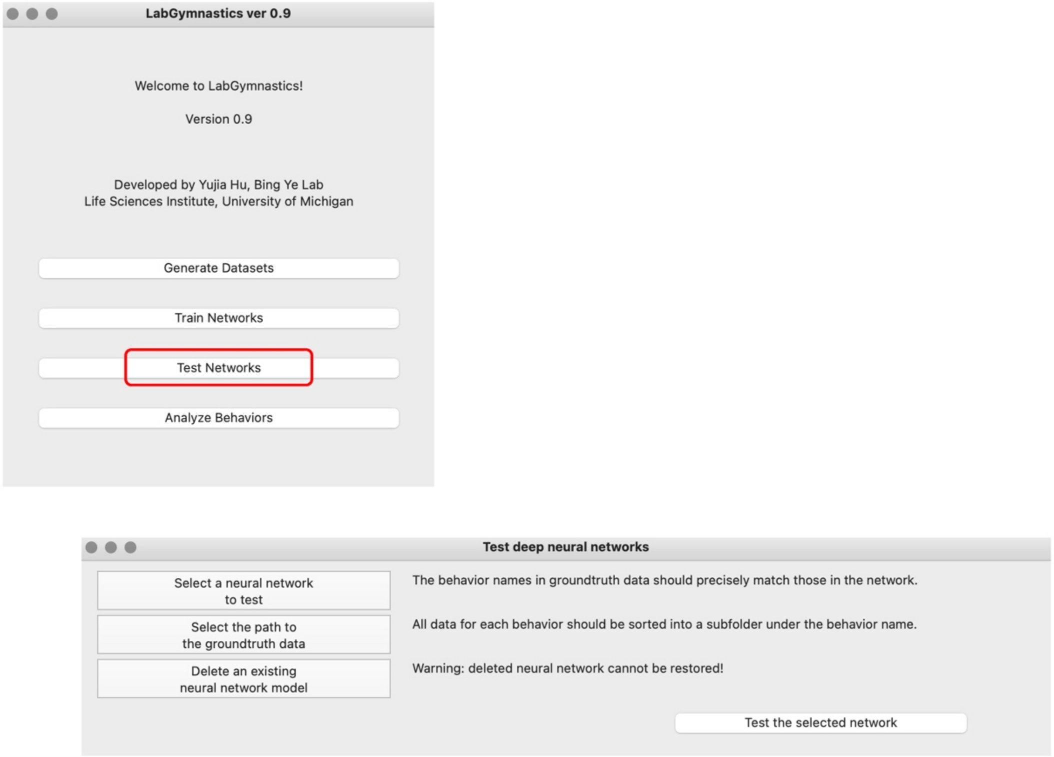


**Supplementary Figure 4. The GUI of Test Networks functional unit.**
This unit offers options for testing a trained Categorizer. Users can the Generate Datasets functional unit to generate behavioral data and sort them into different folders under the behavior names, and then use the sorted ground truth dataset to test the selected Categorizer in unbiased manner.

### Supplementary Figure 5

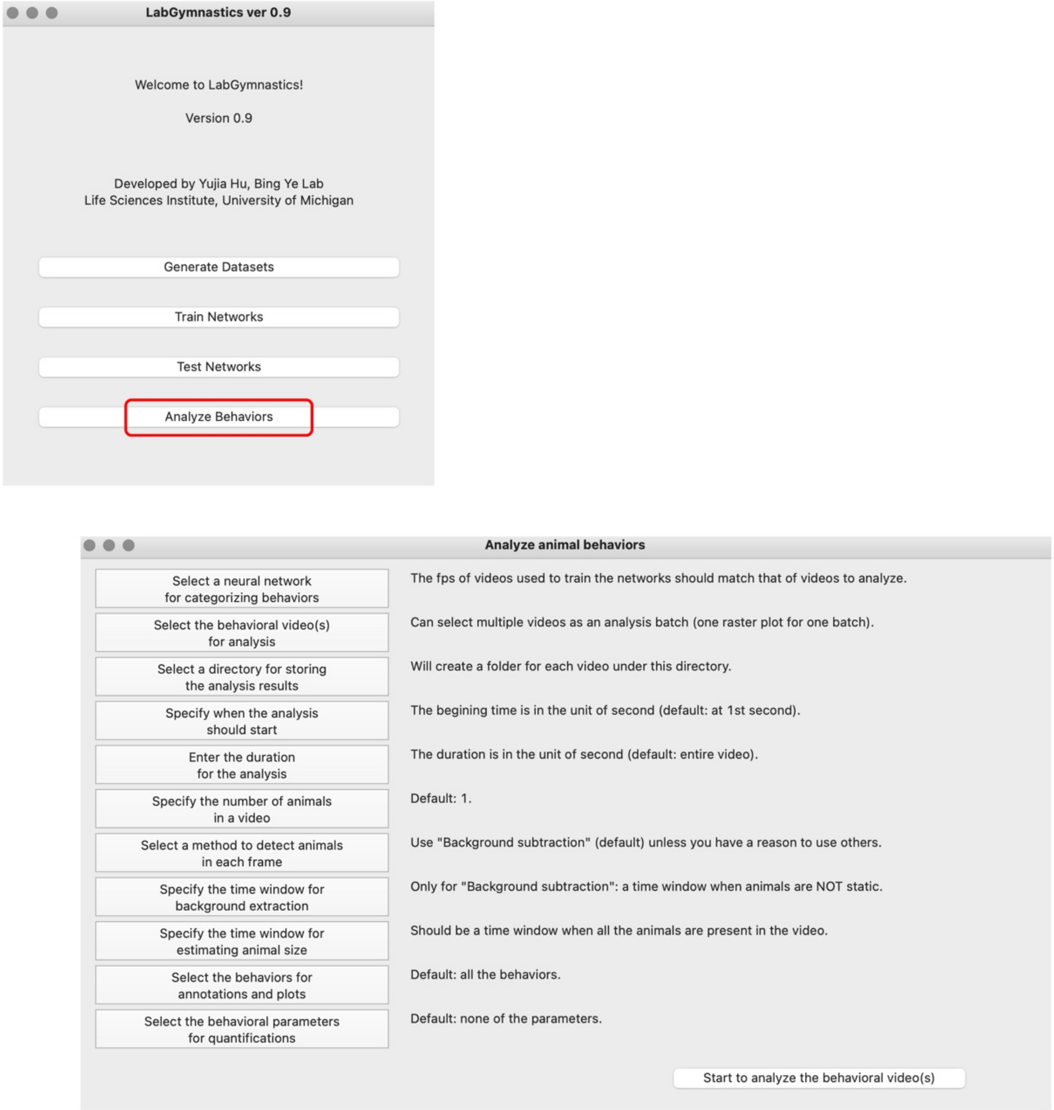


**Supplementary Figure 5. The GUI of Analyzer Behaviors functional unit.** This unit offers options for batch processing of behavioral videos to analyze animal behaviors. For each analyzed behavioral video, this unit outputs a video with annotations of behaviors and individual Microsoft Excel / CSV files for the values of all behavioral parameters for each behavior category and each animal. This unit also outputs a raster plot for all the behavioral events of each animal in one analysis batch. The colors and their intensities in the raster plot, which are user definable, represent the categories and the probabilities for the behaviors, respectively. The functional buttons are on the left side; the short explanations and recommendations are on the right side.

### Supplementary Figure 6

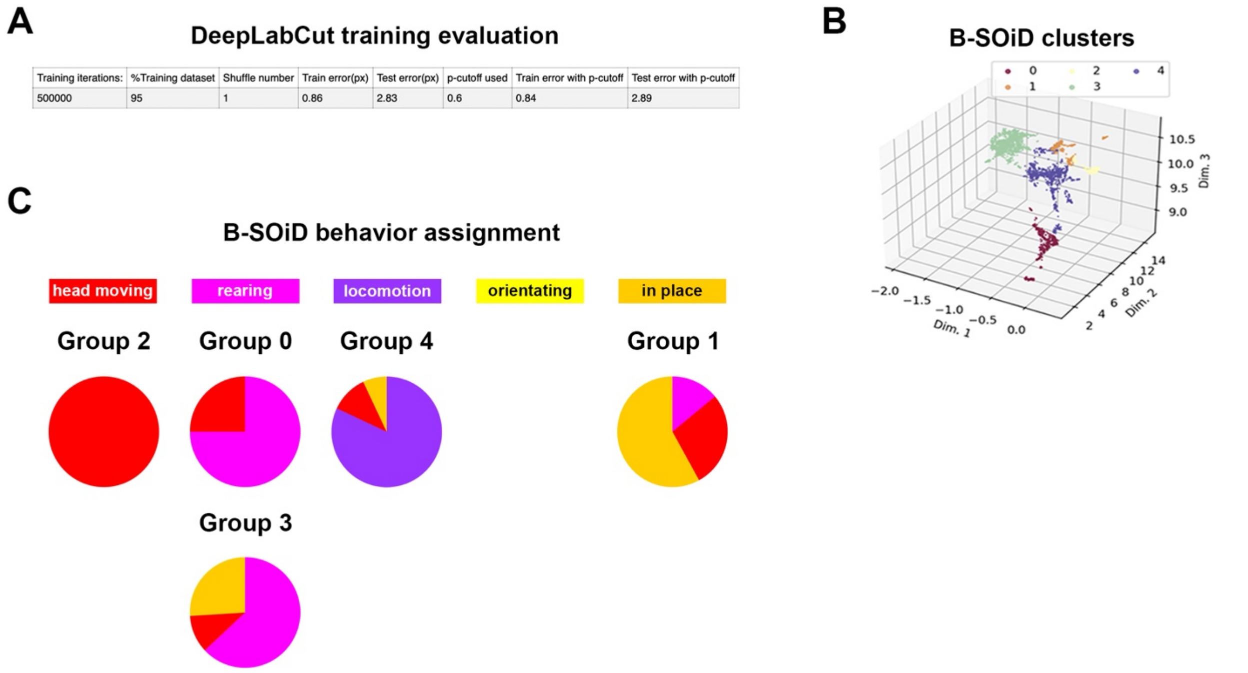


| Training iterations: | %Training dataset | Shuffle number | Train error(px) | Test error(px) | p-cutoff used | Train error with p-cutoff | Test error with p-cutoff |
|---|---|---|---|---|---|---|---|
| 500000 | 95 | 1 | 0.86 | 2.83 | 0.6 | 0.84 | 2.89 |

**Supplementary Figure 6. The training metrics of DLC and B-SOiD in benchmark comparison.**

A. The evaluation of training in DLC on the training dataset.

B. The best fit of clustering (5 groups) in B-SOiD using the outputs from DLC on the training dataset.

C. The behavior assignment of the 5 groups clustered by B-SOiD on the training dataset. All video examples in group 2 showed head movement behavior and group 2 was assigned as 'head movement'; most of the videos examples in group 0 and group 3 showed rearing behavior and group 0 and group 3 were assigned as 'rearing'; most of the video examples in group 4 showed locomotion behavior and group 4 was assigned as 'locomotion'; none of the video examples in all groups showed complete orientating behavior; most of the video examples in group 1 showed in place resting behavior and group 1 was assigned as 'in place'.

| LarvaN (overall accuracy: 0.98) | | | |
|---|---|---|---|
| Animation Analyzer | 8 x 8 x 1, level 1 | | |
| Pattern Recognizer | 32 x 32 x 3, level 2 | | |
| | precision | recall | f1 |
| crawling | 0.99 | 0.97 | 0.98 |
| curling | 0.97 | 0.97 | 0.97 |
| immobile | 0.98 | 1 | 0.99 |
| rolling | 0.97 | 0.98 | 0.97 |
| turning | 0.98 | 0.98 | 0.98 |
| uncoiling | 0.96 | 0.95 | 0.96 |

| Categorizer for Larvae #2 (overall accuracy: 0.93) | | | |
|---|---|---|---|
| Animation Analyzer | 8 x 8 x 1, level 1 | | |
| Pattern Recognizer | 32 x 32 x 3, level 3 | | |
| | precision | recall | f1 |
| crawling | 0.94 | 0.96 | 0.95 |
| curling | 0.92 | 0.88 | 0.9 |
| immobile | 0.97 | 0.95 | 0.96 |
| rolling | 0.94 | 0.95 | 0.95 |
| turning | 0.89 | 0.93 | 0.91 |
| uncoiling | 0.92 | 0.91 | 0.91 |

| Categorizer for Rats #1 (overall accuracy: 0.74) | | | |
|---|---|---|---|
| Animation Analyzer | 16 x 16 x 1, level 2 | | |
| Pattern Recognizer | 32 x 32 x 3, level 3 | | |
| | precision | recall | f1 |
| body grooming | 0.69 | 0.73 | 0.71 |
| face grooming | 0.48 | 0.68 | 0.56 |
| head swaying | 0.66 | 0.74 | 0.7 |
| locomotion | 0.91 | 0.91 | 0.91 |
| orientating | 0.8 | 0.8 | 0.8 |
| rearing | 0.8 | 0.65 | 0.72 |
| in place | 0.58 | 0.56 | 0.57 |
| still | 0.89 | 0.82 | 0.85 |

| Categorizer for Rats #2 (overall accuracy: 0.8) | | | |
|---|---|---|---|
| Animation Analyzer | 32 x 32 x 1, level 4 | | |
| Pattern Recognizer | 64 x 64 x 3, level 4 | | |
| | precision | recall | f1 |
| body grooming | 0.78 | 0.71 | 0.74 |
| face grooming | 0.65 | 0.5 | 0.57 |
| head swaying | 0.74 | 0.76 | 0.75 |
| locomotion | 0.95 | 0.95 | 0.95 |
| orientating | 0.88 | 0.91 | 0.89 |
| rearing | 0.8 | 0.85 | 0.83 |
| in place | 0.69 | 0.65 | 0.67 |
| still | 0.8 | 0.86 | 0.83 |

| Categorizer for Rats #3 (overall accuracy: 0.76) | | | |
|---|---|---|---|
| Animation Analyzer | 32 x 32 x 1, level 5 | | |
| Pattern Recognizer | 64 x 64 x 3, level 5 | | |
| | precision | recall | f1 |
| body grooming | 0.8 | 0.6 | 0.69 |
| face grooming | 0.53 | 0.56 | 0.54 |
| head swaying | 0.72 | 0.79 | 0.76 |
| locomotion | 0.89 | 0.86 | 0.88 |
| orientating | 0.81 | 0.83 | 0.82 |
| rearing | 0.78 | 0.81 | 0.79 |
| in place | 0.62 | 0.61 | 0.61 |
| still | 0.84 | 0.84 | 0.84 |

| RatA (overall accuracy: 0.88) (after refining labeling) | | | |
|---|---|---|---|
| Animation Analyzer | 32 x 32 x 1, level 4 | | |
| Pattern Recognizer | 64 x 64 x 3, level 4 | | |
| | precision | recall | f1 |
| body grooming | 0.89 | 0.88 | 0.88 |
| face grooming | 0.82 | 0.8 | 0.81 |
| head swaying | 0.75 | 0.88 | 0.81 |
| locomotion | 0.95 | 0.98 | 0.96 |
| orientating | 0.96 | 0.92 | 0.94 |
| rearing | 0.88 | 0.88 | 0.88 |
| in place | 0.87 | 0.7 | 0.77 |
| still | 0.89 | 0.93 | 0.91 |

| Categorizer for Mice #1 (overall accuracy: 0.85) | | | |
|---|---|---|---|
| Animation Analyzer | 32 x 32 x 1, level 2 | | |
| Pattern Recognizer | 32 x 32 x 3, level 2 | | |
| | precision | recall | f1 |
| behind the wheel | 0.94 | 0.94 | 0.94 |
| body grooming fv | 0.67 | 0.44 | 0.53 |
| body grooming sv | 0.82 | 0.62 | 0.71 |
| chewing fv | 0.63 | 0.68 | 0.65 |
| chewing sv | 0.74 | 0.85 | 0.79 |
| coming down | 0.89 | 0.99 | 0.94 |
| face grooming fv | 0.66 | 0.83 | 0.74 |
| face grooming sv | 0.86 | 0.67 | 0.75 |
| foraging fv | 0.85 | 0.89 | 0.87 |
| foraging sv | 0.82 | 0.93 | 0.87 |
| rearing up | 0.84 | 0.83 | 0.83 |
| resting on the wheel | 0.91 | 0.92 | 0.92 |
| running on the wheel | 0.93 | 0.93 | 0.93 |
| sniffing fv | 0.59 | 0.57 | 0.58 |
| sniffing sv | 0.87 | 0.74 | 0.8 |
| standing | 0.93 | 0.87 | 0.9 |
| turning front | 0.71 | 0.75 | 0.73 |
| turning side | 0.78 | 0.67 | 0.72 |
| unknown bv | 0.94 | 0.9 | 0.91 |
| walking | 0.93 | 0.9 | 0.91 |

| MouseH (overall accuracy: 0.9) | | | |
|---|---|---|---|
| Animation Analyzer | 64 x 64 x 1, level 4 | | |
| Pattern Recognizer | 64 x 64 x 3, level 4 | | |
| | precision | recall | f1 |
| behind the wheel | 0.94 | 0.91 | 0.93 |
| body grooming fv | 0.71 | 0.83 | 0.77 |
| body grooming sv | 0.85 | 0.79 | 0.82 |
| chewing fv | 0.75 | 0.72 | 0.73 |
| chewing sv | 0.89 | 0.78 | 0.83 |
| coming down | 0.96 | 0.96 | 0.96 |
| face grooming fv | 0.85 | 0.93 | 0.89 |
| face grooming sv | 0.83 | 0.83 | 0.83 |
| foraging fv | 0.93 | 0.92 | 0.93 |
| foraging sv | 0.85 | 0.94 | 0.9 |
| rearing up | 0.92 | 0.92 | 0.92 |
| resting on the wheel | 0.97 | 0.85 | 0.9 |
| running on the wheel | 0.91 | 0.98 | 0.94 |
| sniffing fv | 0.68 | 0.57 | 0.62 |
| sniffing sv | 0.84 | 0.89 | 0.86 |
| standing | 0.94 | 0.95 | 0.94 |
| turning front | 0.76 | 0.81 | 0.79 |
| turning side | 0.92 | 0.79 | 0.85 |
| unknown bv | 0.97 | 0.96 | 0.97 |
| walking | 0.88 | 0.92 | 0.9 |

**Supplementary Table 1. Example showing the training metrics for all tested Categorizers in selecting the ones that are most suitable for each of the 3 behavioral datasets.** We started from the simplest networks for each dataset and gradually increase the network complexity until the training performance was satisfying.

| Building data (data augmentation) | | |
|---|---|---|
| **Augmentation methods** | **The amount of data pair** | **Processing time** |
| Rotation, Flipping, Brighteness change, Deletion | Before: 2,499<br>After: 70,465 | 14 min |
| **Training Categorizer** | | |
| **Settings of Categorizer** | **Training parameters** | **Processing time** |
| LarvaN<br>(Animation Analyzer 1, 8 x 8 x 1;<br>Pattern Recognizer 2, 32 x 32 x 3) | 2,187 steps / epoch, automatically stopped at 24th epoch | ~104 msec / step<br>~230 sec / epoch<br>~ 1.6 hr in total |
| **Analyzing behaviors (LarvaN used for Categorizer)** | | |
| **Video parameters** | **Processing time** | |
| Frame size: 1280 x 720<br>Frame rate: 30 frames / sec<br>The number of animals: 10<br>Duration: 30 sec | Categorizing behaviors: 126 sec<br>Quatifying behavioral parameters: 43 sec | |

**Supplementary Table 2. The typical stepwise processing time in *LabGym*.**

These computational procedures were performed on a MacBook Pro 13 inch (early 2015 model) with Mac OS Monterey (version 12.0.1), 3.1 GHz Dual-Core Intel Core i7 processor, 16 GB 1867 MHz DDR3 memory, and Intel Iris Graphics 6100 1536 MB graphics.

**Supplementary Video 1**: Examples of standard visualizable behavioral datasets generated by *LabGym*

**Supplementary Video 2**: *LabGym* reliably tracks animals in videos with poor contrast and shifting illumination

**Supplementary Video 3**: *LabGym* reliably tracks animals performing non-interactive behaviors

**Supplementary Video 4**: *LabGym* reliably tracks animals performing social behaviors

**Supplementary Video 5**: Larva behavioral dataset: nociceptive behavior subset; mechanosensory behavior subset

**Supplementary Video 6**: Rat behavioral dataset

**Supplementary Video 7**: Mouse behavioral dataset

**Supplementary Video 8**: *LabGym* accurately categorizes user-defined larva behaviors elicited by optogenetic activation of larval nociceptors

**Supplementary Video 9**: *LabGym* accurately categorizes user-defined rat behaviors in tests for psychomotor activities

**Supplementary Video 10**: *LabGym* accurately categorizes user-defined mouse behaviors in a cage equipped with a running wheel

**Supplementary Video 11**: *LabGym* captures the changes in larva nociceptive behaviors upon LK neuron inhibition

**Supplementary Video 12**: *LabGym* captures the differences in larva behaviors elicited by different intensities of sound

**Supplementary Video 13**: *LabGym* captures the sensitization in rat psychomotor activities after repeated amphetamine injections

**Supplementary Video 14**: *LabGym* captures the decline caused by aging in the speed of running on the wheel

**Supplementary Video 15**: *LabGym* accurately identifies rat psychomotor activity in ‘Baseline’ video in benchmark comparison

**Supplementary Video 16**: *LabGym* accurately identifies intensified rat psychomotor activity in ‘Post-treatment’ video in benchmark comparison

**Supplementary Video 17**: *LabGym* generalizes in categorizing rat psychomotor activity in ‘Different enclosure’ video in benchmark comparison
